# Towards transferable explicit-solvent coarse-grained models for biomolecular condensates

**DOI:** 10.64898/2026.08.27.747511

**Authors:** Fran Bačić Toplek, Luis Borges-Araujo, Kresten Lindorff-Larsen, Ralf Everaers, Paulo C. T. Souza, Tatiana I. Morozova

## Abstract

Biomolecular condensates formed by intrinsically disordered proteins require molecular models that accurately describe proteins in both dilute solution and condensed phases. Explicit-solvent coarse-grained models offer an attractive balance between chemical resolution and computational efficiency. Yet, it remains unclear whether improving dilute-state properties is sufficient to obtain an accurate description of condensates. Here, we address this question by introducing minimal modifications to the Martini 3 force field that combine recent advances in bonded interactions with refined protein–water interactions and strengthened glycine self-interactions, while preserving the underlying chemical transferability of the model. The resulting model substantially improves the description of single-chain conformations across a diverse benchmark of disordered proteins. We then investigate phase separation of the well-characterized low-complexity domain of heterogeneous nuclear ribonucleoprotein A1 and its sequence variants. The model reproduces several key physicochemical properties of biomolecular condensates, including chain expansion in the dense phase, sequence-dependent intermolecular contacts, protein diffusion and its relation to single-chain dimensions, and hydration, while revealing quantitative limitations in condensate density, phase equilibria, and ion partitioning. Our results show that improving dilute-state behaviour translates into a better description of condensed-phase properties, including condensate density, but is not sufficient to quantitatively reproduce the equilibrium between the dilute and dense phases.

## Introduction

Biomolecular condensates (BCs) play an essential role in cellular organisation, serving as membraneless compartments that often contain proteins and nucleic acids^1–3^. By concentrating selected molecular components in space and time, they regulate diverse cellular processes, including biochemical reactions, signaling, and stress responses, while protecting biomolecules from unfavourable environments ^4^. Proteins involved in BC formation are often multidomain proteins (MDPs) containing intrinsically disordered regions (IDRs) or intrinsically disordered proteins (IDPs)^3,5–7^. In contrast to folded proteins, IDPs and IDRs do not adopt persistent secondary or tertiary structures. Instead, they populate broad and highly dynamic conformational ensembles^7,8^. Their ability to establish weak, transient, and multivalent interactions underlies the formation, organization, and material properties of BCs. Consequently, physicochemical driving forces behind BC formation, conformational dynamics, and solvent effects within the condensates are essential to their functions. Dysregulation of BC formation and dissolution has been associated with pathological processes, including cancer, neurodegenerative disorders, and infectious diseases^4,9,10^.

Experimental characterization of disordered proteins is challenging because of the heterogeneous nature of their conformational ensembles^11^. Techniques such as small-angle scattering^12–14^ and nuclear magnetic resonance^15^, which probe IDPs in environments close to their native conditions, can capture conformational transitions sampled by these proteins. However, the resulting observables are ensemble-averaged and may reflect contributions from multiple distinct conformational states, making them difficult to interpret from experiments alone. Molecular simulations can therefore provide complementary microscopic insight into the physicochemical principles governing the conformational ensembles and phase behaviour of IDPs, which ultimately underlie their biological functions.

Atomistic molecular dynamics simulations provide a high level of molecular detail and have advanced our understanding of the interactions driving phase separation. Single-chain simulations have identified sequence determinants of chain compaction and its relation to intermolecular interactions^16–20^, whereas simulations of peptide fragments^21,22^ and, more recently, systems containing tens of proteins have begun to characterize condensed phases directly^23–27^. Despite these advances, atomistic simulations remain computationally demanding for studying BCs, which typically contain hundreds to thousands of proteins and span length scales from hundreds of nanometers to micrometers. Furthermore, systematically exploring the effects of sequence, temperature, salt concentration, and other environmental variables remains prohibitively expensive at atomistic resolution.

These limitations have motivated the development of residue-level coarse-grained models, most of which represent proteins using one bead per amino acid in implicit solvent^28–37^. These models have successfully reproduced single-chain conformations and, in some cases, phase-separation observables such as saturation concentrations and estimates of critical temperatures^30,34,38–41^. Owing to their computational efficiency, they make it possible to access large system sizes and long timescales. As a result, the structural and dynamical properties of IDPs at condensate interfaces and within condensate interiors have been investigated at molecular resolution^34,38,39^. In addition, material properties, such as sequence-dependent condensate viscosity, were explored computationally^40,41^.

However, this efficiency is achieved through simplified molecular interactions. In particular, the implicit treatment of solvent neglects hydrodynamic interactions, which have been shown to strongly influence the morphology and coarsening dynamics of soft-matter assemblies, including colloidal gels and, more recently, polyelectrolyte coacervates^42–44^. Consequently, implicit-solvent models may provide only an approximate description of nonequilibrium assembly pathways, condensate coarsening, and rheological properties. Moreover, the absence of explicit solvent and ions precludes a direct description of solvent- and ion-mediated interactions, including salt screening, ion partitioning, and solvent-mediated modulation of protein interactions^24,27,45^.

At a finer level of resolution, the Martini3 force field^46^ provides a middle ground between atomistic simulations and residue-level coarse-grained models. By representing amino-acid backbone and side chains, solvent molecules, and ions explicitly, Martini3 retains substantially greater chemical detail while remaining computationally efficient enough to simulate multichain phase-separated systems^47–52^. Recent developments have extended the Martini3 force field to IDPs^50,53,54^, MDPs^50,51,53,54^, and nucleic acids^55^, making it a promising framework for chemically resolved simulations of biomolecular condensates.

An important concept that aids in rationalizing the formation of BCs is that the conformational properties of isolated IDPs are closely related to their propensity to phase separate, reflecting the similarity between intra- and intermolecular interactions^16,56,57^. This relationship has been extensively validated experimentally and computationally^58–62^ and has motivated the parameterisation of several recent coarse-grained models against experimental single-chain observables^32,61,63^. However, it remains an open question whether improving dilute-state conformations alone is sufficient to obtain a transferable description of biomolecular condensates.

Here, we address this question by first refining the description of IDP single-chain dimensions and testing the transferability of these improvements across a diverse set of IDPs. We then examine how these improvements translate to condensate properties using the lowcomplexity domain of heterogeneous nuclear ribonucleoprotein A1 (A1-LCD). A1-LCD is one of the best-characterized model systems for biomolecular phase separation. It has been extensively investigated experimentally and computationally, with measurements available for dilute- and condensed-phase properties, including saturation concentrations, residue-level interactions, ion partitioning, and viscoelasticity^41,60,61,64–66^. At the same time, the Martini3-IDP model^50,51^ fails to quantitatively reproduce its single-chain dimensions and phase behaviour, making A1-LCD a stringent benchmark for assessing the transferability of explicit-solvent coarse-grained models.

We first introduce minimal modifications to the Martini3-IDP model ^50^ that improve the description of isolated IDPs while preserving the underlying chemical transferability of the Martini3 force field, resulting in a new condensate-oriented model termed Martini3-IDP-C. Specifically, we increase protein backbone–water interactions and introduce a targeted Gly-specific modification that strengthens Gly–Gly interactions. These modifications substantially improve the transferability of the model, yielding more accurate single-chain conformations for a diverse benchmark of intrinsically disordered proteins beyond the A1-LCD family. We then investigate how these improvements translate to the condensed phases through large-scale simulations of A1-LCD condensates. We characterize condensate density, IDP conformations, intermolecular contacts, diffusion, hydration, and ion partitioning. Improved agreement with dilute-state properties translates into a more realistic description of several condensed-phase observables including the potential to capture sequence-dependent differences. At the same time, important challenges remain regarding protein densities in the dilute and dense phases, phase equilibria, and ion partitioning, demonstrating that accurately reproducing dilute-state observables alone is insufficient to quantitatively describe BCs.

Together, our results systematically identify which properties of biomolecular condensates are recovered by improving the description of proteins in dilute solution and which require additional physical ingredients. More broadly, this work proposes a shift in the focus from developing improved parameterisations toward understanding the physical requirements for transferable explicit-solvent coarse-grained modelling of biomolecular condensates.

## Results

### Refining M3-IDP for a diverse set of disordered proteins

We first assess the performance of the Martini3-IDP (M3-IDP) model ^50^ in reproducing experimentally measured radii of gyration, *R*_g_, for the low-complexity domain of heterogeneous nuclear ribonucleoprotein A1 (A1-LCD). The benchmark includes the wild-type sequences (WT and WT+NLS) together with variants in which positively charged (−10R+10K and −6R+6K) or aromatic (−12F+12Y and −7Y+7F) residues were systematically substituted^60^. In Fig. 1a, we find that the M3-IDP model systematically underestimates the radii of gyration of all six A1-LCD sequences, in agreement with previous results reported for the WT sequence^50,51,67^.

**Figure 1:**
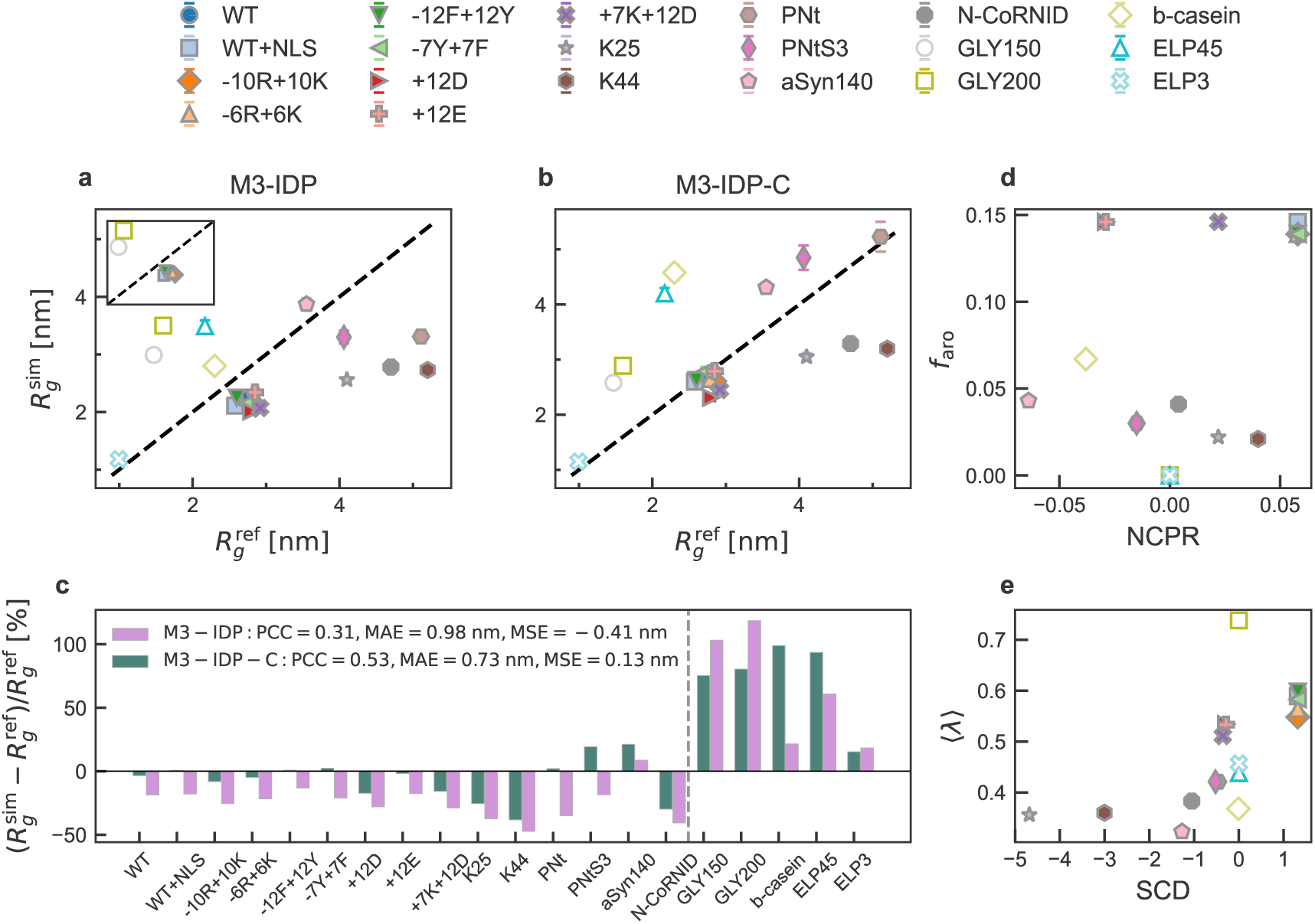
Benchmark of the single-chain size of disordered proteins. **a**, and **b**, Comparison between simulated (sim) and reference (ref) values of the radius of gyration, *R*_g_, for M3-IDP and M3-IDP-C, respectively. The reference values include experimental results (filled symbols) and all-atom simulations (open symbols). The inset in (a) identifies the proteins used for force-field refinement, namely the A1-LCD family variants and poly-Gly. **c**, Relative difference between simulated and reference *R*_g_ values for M3-IDP (purple) and M3-IDP-C (green). Pearson correlation coefficient (PCC), the mean absolute error (MAE), and mean signed error (MSE) are reported in the legend. A vertical dashed line separates experimental and all-atom reference values. **d** and **e**, Physicochemical diversity of selected disordered proteins projected onto net charge per residue (NCPR) *vs* the fraction of aromatic residues, *f*_aro_, and the sequence charge decoration (SCD)^72^ *vs* the average residue stickiness ⟨*λ*⟩^73^, respectively.

Next, we challenge the performance of M3-IDP on a simple sequence from a *naive* polymer-physics perspective—poly-glycine (poly-Gly), which represents the peptide backbone of a protein. As we could not find experimental measurements for sufficiently long poly-Gly chains, we compare our results with all-atom simulations by W. Janke and coworkers^68^, who investigated the coil-to-globule transition of poly-Gly chains of several lengths at room temperature (*T* = 290 K). They found that poly-Gly adopts compact, globular conformations. This behaviour is consistent with previous experimental and computational studies showing that water acts as an effective poor solvent for poly-Gly^69–71^. It reflects the balance between favourable backbone–water interactions and even stronger backbone–backbone interactions, including amide–amide hydrogen bonding and dipolar interactions, which collectively favour chain collapse.

In contrast to all-atom simulations^68^, we find the opposite behaviour in M3-IDP: poly-Gly chains containing 150 and 200 residues remain overly extended, adopting coil-like rather than globular conformations. This observation suggests that the Martini3 force field underestimates the effective self-attraction of the peptide backbone, providing additional motivation for refining both backbone–water and backbone–backbone interactions. Both inconsistencies for A1-LCD sequences and poly-Gly are shown in an inset to Fig. 1a.

In response to our observations that A1-LCD sequences are too compact, while poly-Gly is too extended, we introduce two modifications to the M3-IDP model ^50^. First, we slightly increase the interactions between the backbone (BB) beads and regular-size solvent W beads by increasing the prefactor of the corresponding Lennard-Jones (LJ) potentials by *ε*_VS_*_−_*_W_ = 0.11 kJ/mol. This corresponds to an increase of approximately 2-3 % in the BB-water interactions or 0.04 *k*_B_*T* at T=298 K. Technically, this is realized by adding a virtual site (VS) bead at the position of the BB bead, as suggested in the GoMartini3 approach^74^. This approach is also reminiscent of previous Martini 3 modifications for IDPs and multidomain proteins, where uniform rescaling of protein–water^53^ or protein–protein^54^ interactions—applied either to backbone beads only or to all protein beads—was used to improve agreement with experiments. Second, we reassign the glycine BB type from SP1 to SP1h, thereby strengthening Gly self-interactions. This modification increases self-pair interactions by 0.24 kJ/mol (approximately 8 % or 0.1 *k*_B_*T* at T=298 K). Both modifications are minimal and, importantly, leave all other non-bonded interactions in the M3-IDP force field unchanged, thereby preserving the overall parametrization and chemical transferability of the Martini force field. In what follows, we use the term M3-IDP-C to refer to the M3-IDP model with the two modifications mentioned above, reflecting its intended application to biomolecular condensates.

To test the transferability of the M3-IDP-C model, we simulate (*i*) three additional A1-LCD variants that carry different net charges from those already examined, (*ii*) longer IDP sequences, i.e., with the chain length *N* ranging from 140 to 334 residues, and (*iii*) hydrophobic elastin-like polypeptides (ELPs) and amphiphilic *β*-casein. The latter two sequences are challenging systems to infer single-chain properties in experiments. The former is prone to solidification of initially liquid droplets for sufficiently long sequences^75^, while the latter self-assembles into structures similar to micelles at very low protein concentrations ^14^. Thus, we will again compare our results against recent all-atom simulations^19,20^. The final benchmark comprises twenty IDPs with chain lengths ranging from 18 to 334 residues and reference radii of gyration spanning 1.0 to 5.2 nm. This extension is particularly noteworthy because the M3-IDP model was parameterized and benchmarked primarily using relatively short IDPs containing up to 151 amino acids with experimental radii of gyration below approximately 3.8 nm^50^. It is also consistent with the recent study on single-chain conformations of IDPs, which reports that longer IDPs adopt conformations that are too compact relative to experiment when simulated with the M3-IDP model^67^.

Sequence composition in our benchmark changes considerably as well. Two sequence descriptors that were shown to be correlated with the phase separation (PS) propensity are the net charge per residue (NCPR), which varies from positive to negative, and the fraction of aromatic residues, *f*_aro_, which changes from zero (for poly-Gly and ELPs) to 0.15 for A1-LCD variants shown in Fig. 1d^76^. Other sequence descriptors known to be important for PS are sequence charge decoration (SCD)^40,76,77^, which quantifies how positive and negative charges are distributed along the sequence, and the average residue stickiness ⟨*λ*⟩^73^. Positive SCD values correspond to sequences where charges of equal sign are spaced along the sequence further away than charges of opposite sign, i.e., a well-mixed case, while charge segregation corresponds to negative values. The quantity ⟨*λ*⟩ corresponds to the average relative strength of self-interaction for amino acids *vs* their interaction with water. We project the selected sequences on the (SCD, ⟨*λ*⟩) plane in Fig. 1e. The exact definitions of the sequence descriptors are given in Methods. In conclusion, our benchmark is based on twenty sequences including neutral sequences (poly-Gly, ELPs), sequences with high charge segregation (SCD*<* 0), e.g., K25 and K44, and a well-mixed A1-LCD family.

In Fig. 1a and b, we show the performance of M3-IDP and M3-IDP-C models in reproducing reference *R*_g_ values, respectively. The modification we propose to M3-IDP substantially improves overall model performance by increasing the Pearson correlation coefficient (PCC) between simulated and reference *R*_g_ values and reducing the mean absolute error (MAE), as shown in Fig. 1c. For the complete A1-LCD family, including both refinement and validation sequences, the MAE decreases from 0.60 nm with M3-IDP to 0.18 nm with M3-IDP-C, corresponding to an approximately threefold reduction in error. However, we note that the model performance for *β*-casein and ELP45 worsens. For these long sequences, i.e., *N* = 228 and 209 residues, respectively, the reference data originate from all-atom simulations. Although advanced sampling methods were used in both cases to sample conformational space, and good agreement between the experimental and simulated SAXS profiles was observed for *β*-casein, equilibrating such long IDPs remains challenging. When we exclude all-atom simulations from the reference dataset, the PCC values for M3-IDP and M3-IDP-C increase to 0.65 and 0.68, respectively, while the MAE decreases to 0.95 nm and 0.52 nm, respectively.

### Transfer of M3-IDP-C accuracy from single chains to biomolecular condensates

An accurate representation of single-chain conformations is expected to be an important prerequisite for faithfully reproducing phase equilibria involving biomolecular condensates. To examine whether these improvements are also *sufficient*, we test whether the M3-IDP-C model, which reproduces the radii of gyration of A1-LCD and its variants, also provides an improved description of their condensed phases. The A1-LCD family is well studied both experimentally and using other coarse-grained models, making it an excellent case to probe the accuracy of the M3-IDP-C model^30,41,60,61,64–66^. In particular, we investigate A1-LCD wild type (WT), a sequence where all tyrosine (Tyr, Y) residues are substituted with phenylalanine (Phe, F) −7Y+7F, and *vice versa* −12F+12Y, probing the impact of aromatic residues on A1-LCD propensity to PS. All three sequences carry a net positive charge of +8 *q_e_*. We also include the +12E variant, which carries a net negative charge of −4*q_e_* and contains 12 glutamate (Glu, E) substitutions replacing glycine or serine residues. The latter we simulate at 277 K, while simulations of the WT, −7Y+7F, and −12F+12Y are performed at 293 K to evaluate protein densities at the corresponding experimental conditions.

To reduce finite-size effects in our condensate simulations, we adopt the number of chains and box dimensions recommended in previous coarse-grained simulations of A1-LCD condensates^41^. Specifically, we randomly insert 200 copies of a protein in a rectangular box with the initial dimensions of *L_x_*= *L_y_*= 17 nm and *L_z_*= 120 nm, followed by the addition of solvent and ion beads. In particular, this choice of lateral box dimensions accommodates the broad distribution of protein conformations observed in the condensate, with characteristic *R*_g_ values ranging from approximately 2 to 6 nm (see Fig. 5a), thereby reducing potential interactions with their periodic images. We conduct simulations for at least thirty microseconds, where we seed the dense phase by applying flat-bottomed position constraints to proteins. See Methods and Fig. S2,3 in the Supplementary Information (SI) for further details of the equilibration procedure.

To evaluate the convergence of the slab simulations, we compute the temporal evolution of the mass density profiles of protein, water, and ion beads shown in Fig. S5 in the SI, which no longer change substantially after 15 microseconds. Consequently, we use the last ten microseconds for the analysis of equilibrium properties. In the case of the +12E sequence, which we simulate at a lower temperature (T=277 K), we increase the length of the production run to 60 microseconds and perform analysis over the last 20 microseconds to account for the slower relaxation dynamics of the chains.

When the protein-rich phase is simulated with either version of the M3-IDP model^50,51^, the observed protein density (≈ 692 mg/ml) is almost double that of the corresponding experimental value of approximately 366 mg/ml ^60^. In Fig. 2, we plot the mean protein (mass) density profiles along the *z*-axis for the A1-LCD family obtained with the M3-IDP-C model. While significantly improved (for the WT, the protein density in the slab decreases to 570 mg/ml), the observed protein densities are still 55 % greater than the experimental values.

**Figure 2:**
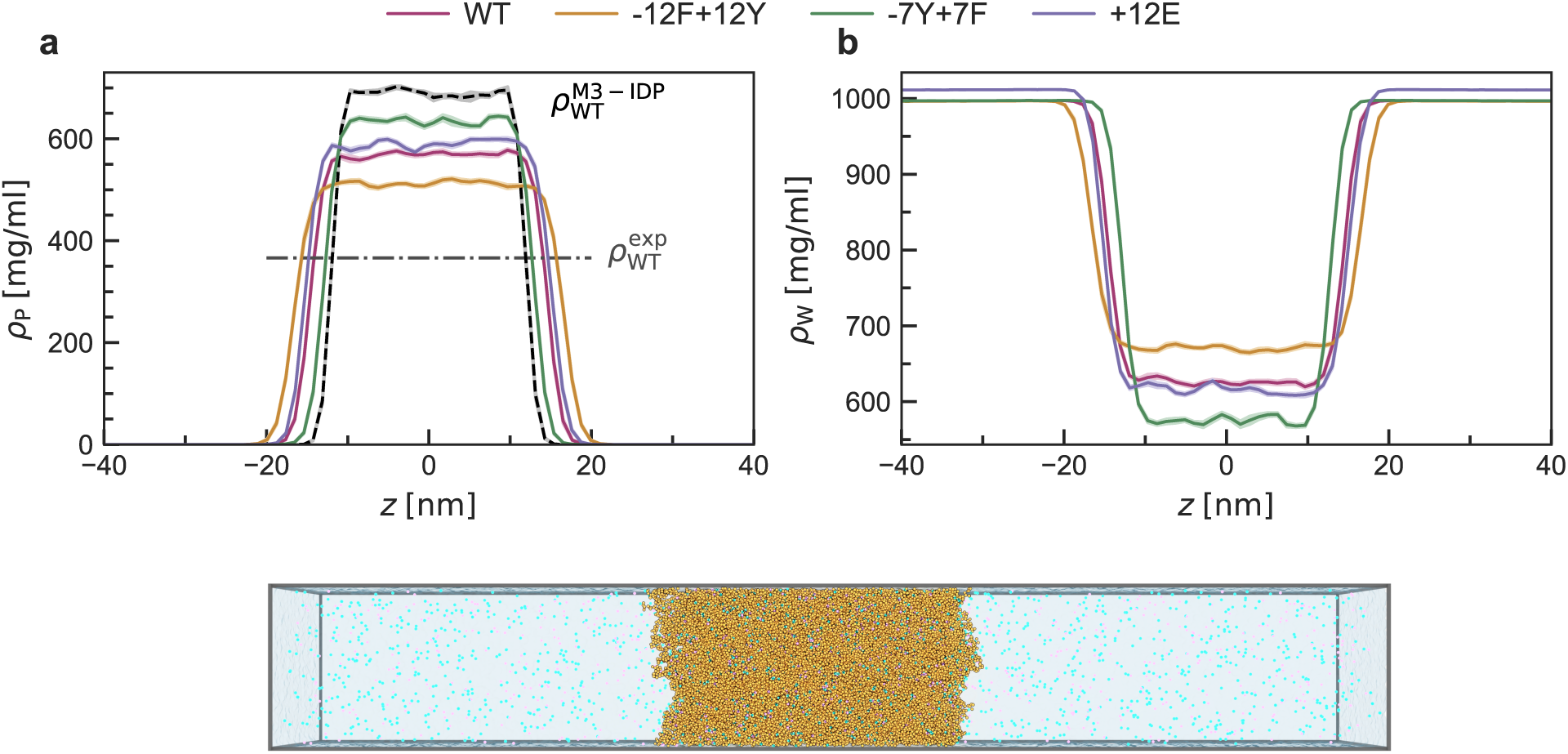
Mean mass density profiles in slab simulations of four A1-LCD variants: **a**, protein (P), where the experimental density of the dense protein phase for WT is shown by black dash-dotted line; **b**, water (W) along *z*-axis. The mean profiles are computed by averaging over one-microsecond window for simulation time between twenty (forty) and thirty (sixty) microseconds for WT (+12E), −12F+12Y, and −7Y+7F. Shaded areas represent the error bars, calculated as the standard error of the mean. A representative snapshot of the WT system is shown below the subplots. Proteins are rendered in yellow, Na^+^ in blue, Cl*^−^* in pink, and water as a light-blue background.

We note that no stable dilute phase was observed for any of the sequences studied here. To verify that this was not an artifact of averaging the density profiles, we analysed the positions of individual protein chains relative to the condensate interface and found no proteins residing in the dilute phase throughout the equilibrated trajectories (see SI for details). We further tested whether the absence of proteins in the dilute phase resulted from the slab initialization by introducing ten additional proteins into the dilute region surrounding an equilibrated WT condensate. These proteins either joined the condensate or formed a transient aggregate that would ultimately reduce the total interfacial free energy by integrating into the slab (Fig. S4). Together, these controls demonstrate that the absence of a dilute phase is not an artifact of the analysis or simulation setup, but rather indicates that the effective protein–protein interactions in the current model remain somewhat too strong. Similar behaviour was reported for the low-sequence-complexity domain of fused in sarcoma protein simulated with the Martini 3 force field^46^, where the unmodified force field produced complete condensation and modest strengthening of protein-water interactions was required to observe the dilute phase. Further refinement of the force field will therefore be required to accurately reproduce dilute-phase protein concentrations and phase equilibria.

Next, we find that the density of the protein-rich phase depends on the protein sequence, with −12F+12Y and −7Y+7F forming the least and most dense condensates, respectively. Experimentally, protein-rich phase densities have been reported to lie within a relatively narrow range of 300 mg/ml for −7Y+7F to 367 mg/ml for WT at *T* = 293 K^60^. In contrast, the densities of the dilute phase vary by approximately a factor of three, with −12F+12Y and −7Y+7F exhibiting the lowest and highest values, respectively^60^. Thus, in experiments, the −12F+12Y sequence displays stronger self-interactions, whereas our simulations show the opposite trend: −7Y+7F exhibits stronger self-association propensity, as reflected in the higher density of the protein-rich phase. We also find that the dense phase is well hydrated, as shown in Fig. 2b, similar to all-atom simulations of biomolecular condensates^21,26^.

Unlike residue-based implicit-solvent coarse-grained models, the Martini3 force field represents both ions and solvent explicitly. Thus, in Fig.3 we investigate charge distributions in the systems by resolving charge density profiles along the slab normal. The temporal evolution of these profiles is provided in the SI (Fig.S7). Since no proteins are found in the dilute phase in the present simulations, the protein charges are concentrated within the condensed phase (Fig.3a). In addition, both Na^+^ and Cl*^−^* exhibit enhanced concentrations inside the condensates relative to the dilute phase for all sequences, although the degree of enrichment depends on sequence composition and net charge (Fig.3b,c). These results are in qualitative agreement with recent measurements of Na^+^ and Cl*^−^* activities in A1-LCD condensates by Posey et al.^65^, which showed that positively and negatively charged condensates preferentially partition counterions. However, unlike the experiments, our simulations predict enrichment of both ion species within the condensed phase. Experimentally, positively charged condensates preferentially accumulate Cl*^−^* while excluding Na^+^, whereas the opposite behaviour is observed for negatively charged variants, leading to the description of biomolecular condensates as mesoscale capacitors that store electrical charge through differential ion partitioning across the condensate interface ^65^.

**Figure 3:**
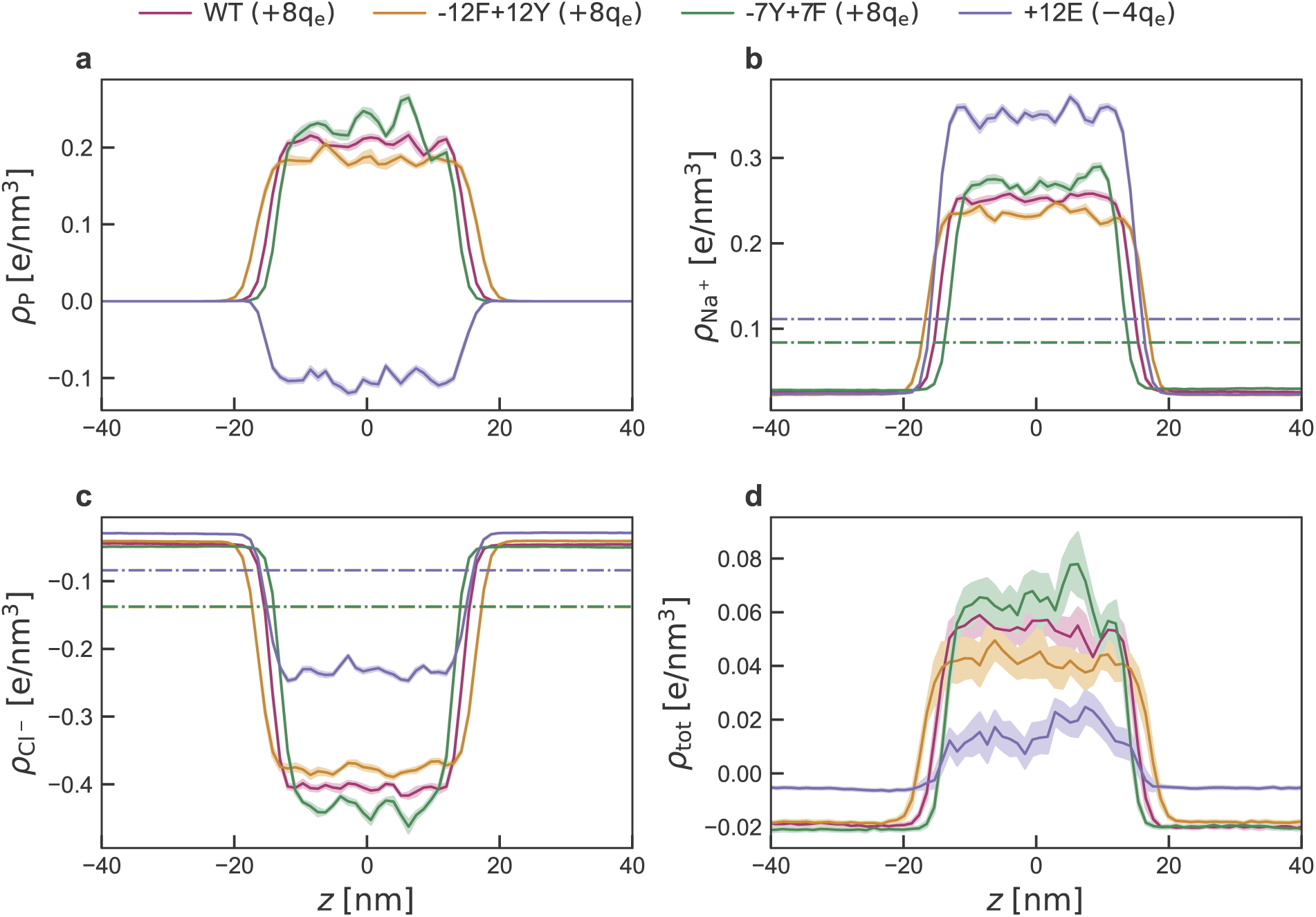
Mean charge density profiles of four A1-LCD variants: **a**, protein (P); **b**, sodium cation (Na^+^); **c**, chloride anion (Cl*^−^*); **d**, the total charge (tot) along *z*-axis. The mean profiles are computed by averaging over one-microsecond window for simulation time between twenty (forty) and thirty (sixty) microseconds for WT (+12E), −12F+12Y, and - 7Y+7F. Shaded areas represent the error bars, calculated as the standard error of the mean. Dash-dotted lines correspond to the bulk ionic densities.

Several factors may contribute to this discrepancy between our numerical results and experiments. In the Martini force field, electrostatic interactions between charged beads are described by Coulomb interactions in a homogeneous dielectric medium with relative permittivity *ε_r_* = 15, whereas water beads are electrically neutral and interact only through Lennard-Jones potentials. The relatively low value of the dielectric constant corresponds to a Bjerrum length, the distance at which the Coulomb interaction between two elementary charges equals the thermal energy, of approximately 3.8−4.0 nm under the present conditions, substantially larger than in water (≈ 0.7 nm), indicating strong short-range electrostatic interactions between charged beads. In contrast, the Debye screening length, the characteristic distance over which electrostatic interactions are screened by mobile ions, estimated from the local ion concentrations shown in Fig.3a-c, is approximately 0.5−0.6 nm in the dilute phase and decreases to about 0.2 nm inside the condensates because of the increased ionic strength. Consequently, electrostatic interactions are energetically strong at contact but screened over subnanometre distances comparable to the size of Martini ion beads. This regime favours local ion pairing and ion–protein associations rather than long-range electrostatic correlations, which may contribute to the simultaneous enrichment of both Na^+^ and Cl*^−^* within the condensates. Additional differences may arise from the coarse-grained description of water, which lacks explicit molecular polarization, and from the finite simulation box, whereas experiments are effectively coupled to an infinite salt reservoir. Lastly, while monovalent ion–water interactions were parameterized in the Martini 3 force field and validated in ion–membrane systems and ionic liquids, protein–ion interactions were less extensively tested, leaving potential room for improvement in the corresponding pair interactions^46^.

Finally, despite the different net charges of the protein sequences, all condensates exhibit a small positive charge density (Fig. 3d), indicating incomplete local electroneutrality. This behaviour is qualitatively consistent with experimental observations that biomolecular condensates behave as weak charge reservoirs^65^.

### M3-IDP-C predicts the key contacts responsible for the condensate formation

Next, we look into key interactions that drive the PS of the A1-LCD family by computing inter-chain contacts. To accurately capture the contact formation between proteins, we adopt the bead-size cut-off distance earlier proposed by Feito et al^41^. Furthermore, we distinguish contributions into contact formation types, i.e., backbone-backbone, backbone-side chain, and side chain-side chain interactions. For technical details of these calculations, see the Methods section.

In Fig. 4a, we show the contact map for the slab formed by the WT sequence using all beads. We observe that the most frequent contacts are formed by residues in the second half of the sequence and often involve Phe residues, irrespective of their position in the sequence. We identify strong contact regions between residues 90–117 and 130–137, which contain Phe, Tyr, Lys, and Arg, in agreement with earlier experimental and numerical observations^41,60^. In addition, we find that methionine at position 93 (Met93) often forms contact pairs with Phe residues. Owing to the explicit representation of side chain and backbone beads in the Martini force field, we can conclude that these interactions are primarily mediated by side chain–side chain contacts, as shown in Fig. 4b–d. For aromatic–aromatic interactions, the geometrical resolution of the Martini 3 force field allows different relative orientations of aromatic groups, including stacked and T-shaped configurations, to be distinguished to some extent, although their relative stability may not always be quantitatively reproduced^78^. Cation–*π* interactions, on the other hand, are not explicitly encoded but can be effectively captured through the nonbonded interactions between charged and aromatic beads. Thus, the preferential Phe–Phe and Phe–Arg side-chain contacts observed here are consistent with aromatic–aromatic and cation–*π*-like interactions, respectively. We note that in Fig. 4d, the dark blue regions corresponding to strictly zero contact frequency arise from Gly residues, which are modeled by a backbone bead only. The backbone-side chain contribution shows a similar pattern but is less frequent than the side chain-side chain contribution. Often, when two side chains of a residue pair are in contact, their backbone beads are also in proximity. In contrast, backbone-backbone interactions are rare; see Fig. 4b. We find similar results for the other three sequences, shown in the SI, Fig. S8-10.

**Figure 4:**
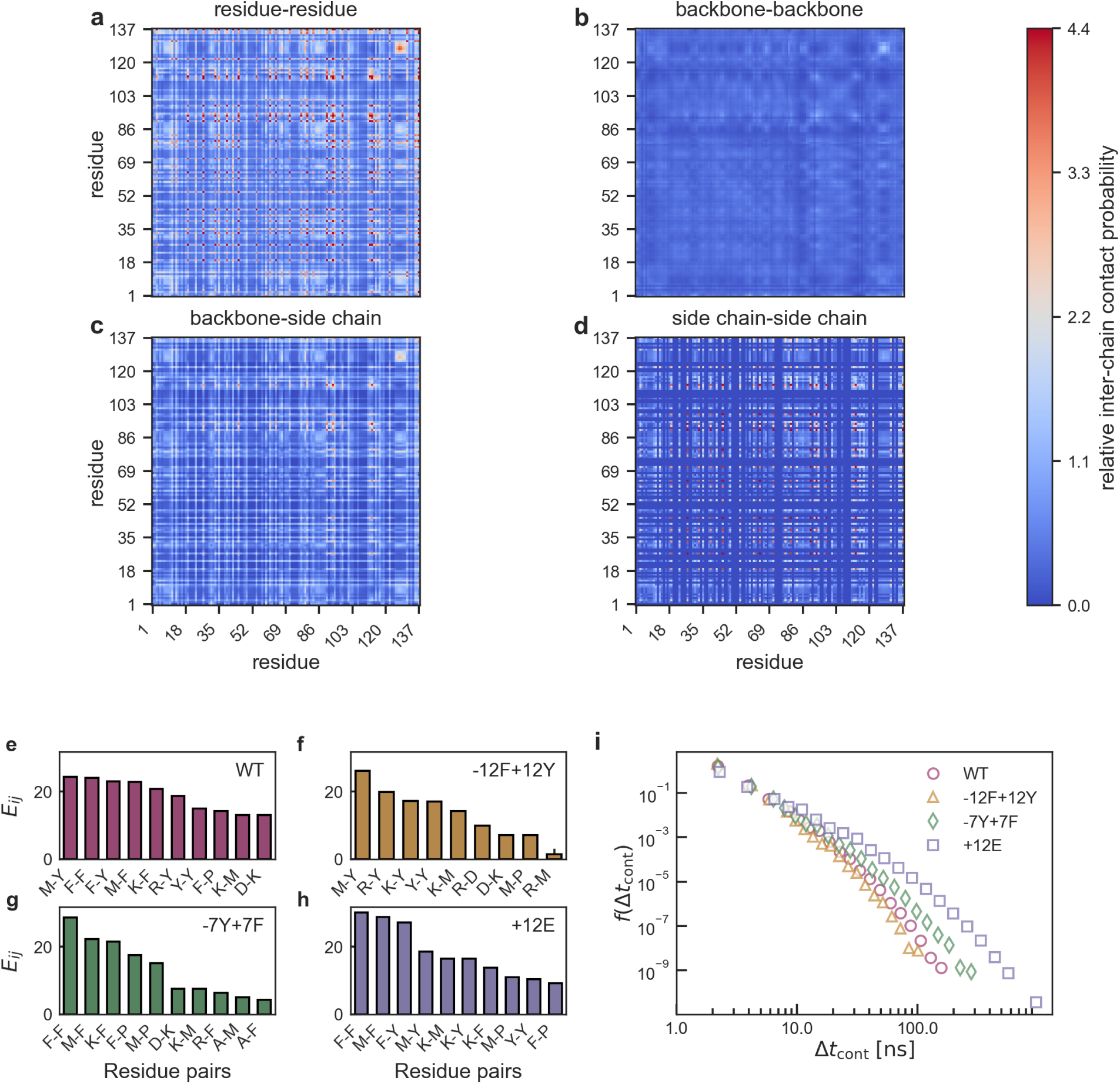
Contact formation in A1-LCD condensates. **a**-**d**, Inter-chain contact probability for the WT sequence using all, only backbone, backbone and side chain, and side chain beads. The absolute value is normalized by the number of possible chain pairs and the number of snapshots in a trajectory, and scaled by the mean value of the residue-residue contacts. The maximum value is capped at the 99th percentile of the relative residue-residue contacts. **e**-**h**, Enrichment of inter-chain contact pairs *E_ij_* for WT,-12F+12Y, −7Y+7F and +12E sequences. **i**, Probability density function of the contact lifetime.

Furthermore, we identify residue pairs that preferentially contribute to interchain contacts irrespective of their position along the sequence, as shown in Fig. 4e–h. We compute a composition-normalized enrichment *E_ij_* of significant residue-type *i* − *j* contact pairs, with values *E_ij_ >* 1 indicating preferential contact formation relative to sequence composition (see Methods for technical details on this calculation). For all four sequences, the most frequently occurring significant contacts are formed by F/Y–F/Y, M–F/Y, and K/R–F/Y pairs. While *π*–*π*, sulfur–*π*, and cation–*π* interactions are not explicitly encoded in the Martini3 potential, they can be effectively represented through nonbonded interactions between the corresponding beads, and the observed contacts are consistent with residue-pair preferences associated with these interactions.

For sequences containing both phenylalanine and tyrosine residues, we find that F-F contacts are more prominent than F-Y contacts, followed by Y-Y contacts (see Fig. 4e, h). Naturally, the relative preference for phenylalanine (F) or tyrosine (Y) changes when only one of these two amino acids is available, as shown in Fig. 4f *vs.* Fig. 4g. Finding this hierarchy of contacts in our simulations can explain the variation in the protein density in the slab shown in Fig. 2a as F content on the sequence increases, i.e., the density *ρ*_P_(−12F+12Y) *< ρ*_P_(WT) *< ρ*_P_(−7Y+7F) suggesting stronger F-F over Y-Y interactions. This trend contrasts with experimental findings, where lower dilute-phase protein densities were reported for the Y-rich sequence, i.e., −12F+12Y, compared with WT and −7Y+7F, indicating stronger protein-protein interactions for this sequence. Accordingly, the following relative interaction strengths were established experimentally: Y-Y*>*Y-F*>*F-F^32,60,61^. The underestimation of Y–Y interactions by the Martini force field is, in part, expected, as a similar tendency was previously observed for its side-chain analogue^78^.

Other important driving forces for the contact formation in our simulations are positively charged-aromatic residue pairs, i.e. R/K-Y/F, followed by electrostatic (D-R/K) and hydrophobic contacts between aromatic residues and proline. This finding is consistent with results obtained for the WT sequence using the one-bead-per-residue CG model CALVADOS 2^41^. Our analysis also ranks M-F/Y contacts among the top ones. Other coarse-grained models that identify M-Y/F interactions as important for WT condensates include HPS^29^, HPS-Urry^31^ and CALVADOS 2^34^. Overall, it is remarkable that M3-IDP-C captures the main contact patterns associated with A1-LCD condensate formation, despite not being explicitly parameterized against these interactions, demonstrating that the Martini building-block approach that relies on partitioning behaviour of small molecules can be transferable across different systems and contexts.

To complete the picture of residue contacts in A1-LCD sequences, we also examine into contact formation in single-protein systems, i.e., intra-chain contacts. We plot the most significant intra-chain contact pairs in Fig. S11 in the SI. As for condensates, we find that F/Y contacts are among the most dominant ones; the strength of R-Y/F and electrostatic interactions, e.g., R-D, is increased compared to multi-chain systems. Also, contacts between Met and aromatic residues are not as prominent as in protein condensates, while interactions between polar amino acids Ser, Asn, Gln, and negatively charged Asp make up a substantial fraction of intra-chain contacts. Thus, our results support the expected similarity between intrachain contacts formed by isolated proteins in solution and interchain contacts formed in the dense phase^32,58–61^.

Finally, we examine the timescales of contact formation. The probability density function (PDF) of contact lifetimes, *f* (Δ*t*_cont_), shown in Fig. 4i, is dominated by short-lived contacts, i.e., ≤ 10 ns, highlighting their transient nature. Since the +12E sequence was simulated at a lower temperature, its dynamics, including contact lifetimes, are slower than those of the other sequences. Comparing contact lifetimes among the remaining three sequences, we observe a higher probability of long-lived contacts for the F-rich sequence, −7Y+7F, followed by WT and −12F+12Y. This again suggests stronger interaction propensities for Phe compared with Tyr residues in the M3-IDP-C model.

In summary, within the Martini force-field representation, intermolecular contacts are predominantly associated with aromatic residues, particularly through aromatic–aromatic pairs involving Phe/Tyr residues and positively charged–aromatic pairs, in agreement with previous experimental and numerical findings. In addition, methionine emerges as an important contributor to intermolecular association through strong Met–aromatic contacts^18^ in multi-chain systems, whereas electrostatic interactions appear to play a more important role in solution.

### Disordered proteins are more extended when forming condensates

Next, we investigate protein conformations at the condensate interface and in the condensate interior and compare them with the results obtained from single-chain (SC) simulations, corresponding to the infinite-dilution case. We use the protein’s COM position to assign the protein to the corresponding region within a condensate. Specifically, we classify protein *i* to be part of the bulk region (cond. bulk) if the *z*-coordinate of its COM satisfies: −*z*_DS_ + *d <* 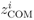 *< z*_DS_ − *d*, and to the interfacial region (cond. interface) otherwise, see the inset in Fig. 5a. The values of *z*_DS_ and *d* are the position of the dividing surface and the interfacial thickness, respectively, extracted from the fits with a hyperbolic tangent function to the average protein mass density profiles shown in Fig 2a (see SI for the details).

**Figure 5:**
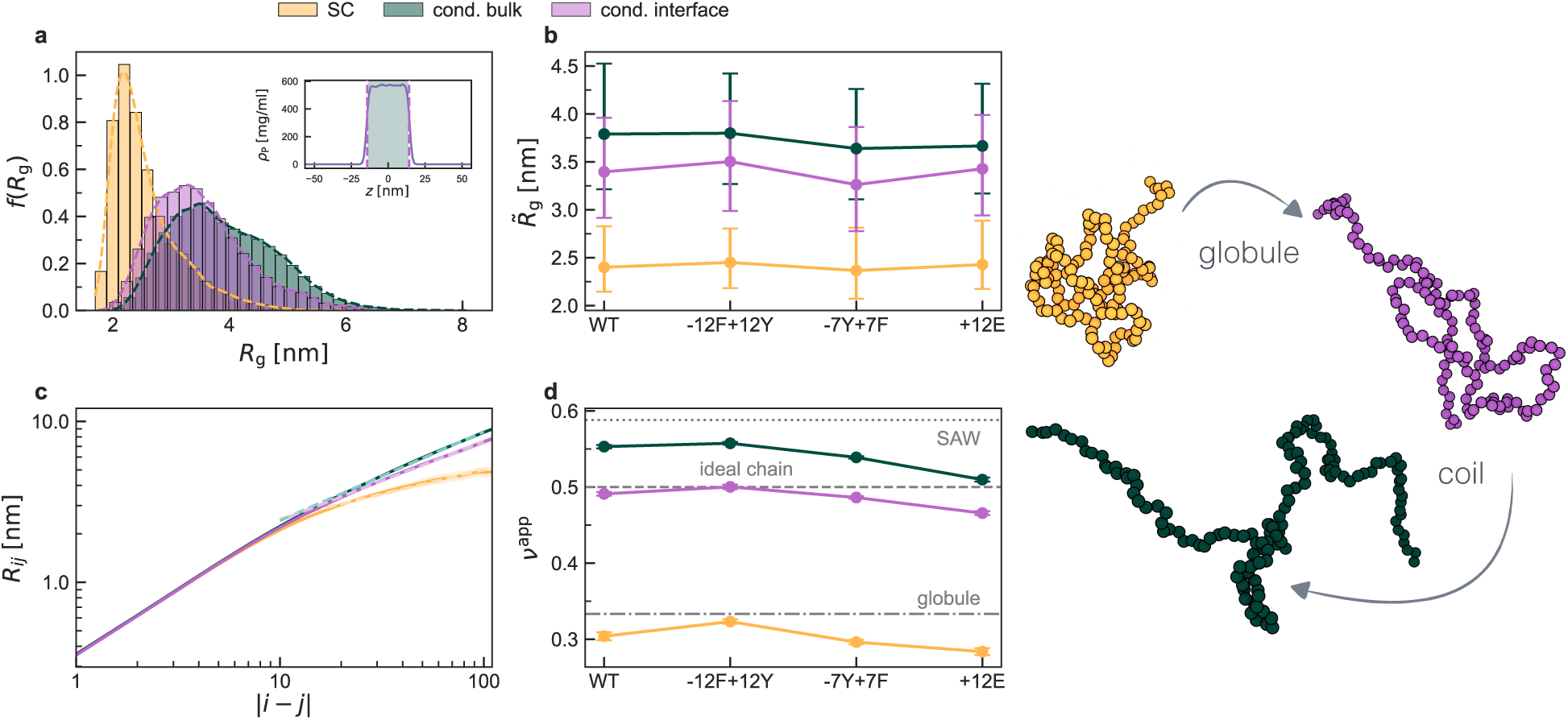
Protein conformations in solution and in condensates: **a**, Probability density function (PDF) of the radius of gyration,*R*_g_, for the WT sequence in the dilute (single-chain, SC) limit, at the interface (cond. interface), and inside the protein-rich phase (cond. bulk); Inset: mean mass density profile for WT. Vertical dashed lines indicate the condensate boundaries, while the green shaded region represents the condensate interior. **b**, Median of the PDF of the radius of gyration *R̃*_g_ for A1-LCD sequences, where bars represent the interquartile range (Q1–Q3); **c**, root-mean-square distances *R_ij_* between residues *i* and *j* as a function of the distance along the sequence |*i* − *j*| for WT sequence. Dashed curves correspond to the fits with the expression *R_ij_* = *R*_0_ |*i* − *j*|*^ν^*^app^ over the separation distances 10 ≤ |*i*−*j*| ≤ 100, where *R*_0_ is the scaling prefactor, and *ν*^app^ is the apparent Flory exponent; **d**, the apparent Flory exponent for A1-LCD sequences. Dotted, dashed, and dash-dotted lines correspond to the self-avoiding walk (SAW), *ν* = 0.588, ideal chain, *ν* = 1*/*2, and globule, *ν* = 1*/*3, limits, respectively. All subplots share the same legend.

In Fig. 5a, we compare PDFs of the radius of gyration, *f* (*R*_g_), for the WT sequence under the dilute and protein-rich conditions. Similar curves are obtained for other A1-LCD variants shown in the SI, Fig. S12. In Fig. 5b, we report the median *R̃*_g_ of *f* (*R*_g_) for all sequences studied, with the interquartile range (*Q*_1_–*Q*_3_) indicating the spread of the distributions. We find that proteins considerably swell when partitioned into condensates with a gradual transition over the interfacial region. This is not surprising, as the aqueous environment is a poor solvent for the A1-LCD family, which is consistent with their PS behaviour under the simulated conditions, while the protein-rich phase resembles ideal solvent conditions. To confirm this reasoning, we compute the root-mean-square internal distances, *R_ij_*, between residues as a function of their separation |*i* − *j*| along the sequence, as shown in Fig. 5c for the WT and Fig. S13 for other sequences. At small separation distances |*i* − *j*| *<* 10, the curves are indistinguishable as imposed by chain connectivity. However, at larger separations along the sequence, the scaling of *R_ij_* ∝ *R*_0_|*i* − *j*|*^ν^*^app^ is qualitatively different for SC and condensed phase conditions, where *R*_0_ is the scaling prefactor, and *ν*^app^ is the apparent Flory exponent^79^. Similar curves are obtained for other A1-LCD variants shown in the SI, Fig. S13. In Fig. 5d, we summarize *ν*^app^ values in dilute and protein-rich conditions. Although both *R*_0_ and *ν*^app^ determine the internal-distance profile, *R*_0_ serves as a fitting parameter; therefore, we focus on *ν*^app^ as the parameter that captures differences in long-range chain scaling. Indeed, under infinite dilution, chains sample compact, globule-like conformations with *ν*^app^ values even below the *ν* ≈ 1*/*3 scaling expected for infinitely long homopolymers under poor-solvent conditions^79^. This apparent discrepancy reflects the finite length and heterogeneous sequence composition of A1-LCD, for which *ν*^app^ should be regarded as an effective descriptor of chain statistics rather than a universal Flory exponent. In contrast, proteins sample more extended conformations at the interface, reaching the scaling for an ideal chain with *ν* ≈ 1*/*2. Proteins are even more extended inside the condensates, with *ν*^app^ being between the ideal chain and self-avoiding walk limits. These findings are consistent with other numerical works that probed conformations of disordered proteins at the interface and inside condensates using all-atom^21^ and other CG models^34,38–40,76,80^. We note that other averaging approaches can produce different protein conformations at the condensate interface^64,81^.

### M3-IDP-C captures protein diffusion within experimental ranges

Finally, we examine the dynamics of the A1-LCD proteins. We first investigate chain dynamics at the condensate interface and in the condensate interior using the classification based on the *z*-coordinate of its COM described in the previous section. In Fig. 6a, we compute PDFs of the protein residence time, *τ_R_*, in the interfacial and bulk regions for WT. Both PDFs are dominated by short residence times, highlighting frequent chain transitions between the interfacial and bulk regions. We note a nonlinear feature in the bulk residence-time distribution at the longest residence time, i.e., 10 *µ*s. This point corresponds to proteins that do not leave the bulk region during the part of the trajectory used for the analysis and therefore remain part of the condensate interior. This population comprises approximately 75% of the chains. Similar behaviour is observed for the other sequences, as shown in Fig. S14.

**Figure 6:**
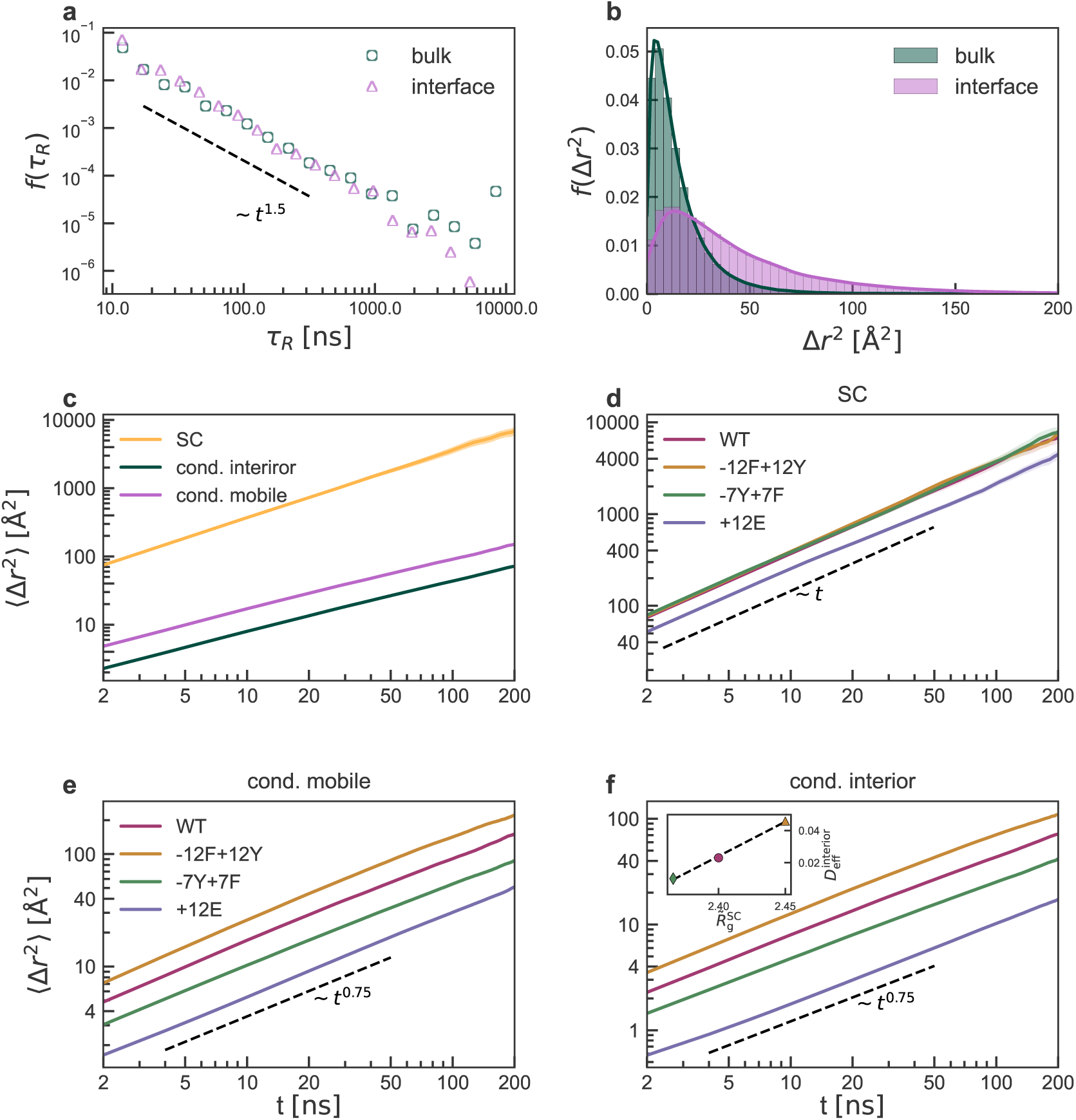
Protein dynamics in solution and in condensates. **a** and **b**, PDFs of the residence time and the squared displacement, Δ*r*^2^, for Δ*t* =20 ns of WT proteins at the interface and inside the condensate. **c**, the mean squared displacement (MSD) ⟨Δ*r*^2^⟩ of WT in solution (single-chain, SC), in the interior (cond. interior), and mobile (cond. mobile) regions of a condensate. **d**-**f**, MSD grouped by conditions, i.e., in solution, mobile, and interior regions of a condensate to highlight sequence specificity. Inset **f**: the correlation between the effective diffusion coefficient computed for the condensate interior, 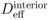, and the median of the radius of gyration for SC, 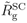. Subplots **d**-**f** share the same legend.

Following the analysis proposed by Yamamoto et al. ^39^, who investigated protein dynamics in IDP condensates using HPS-family models, we fit both PDFs with a power-law distribution with an exponential cutoff, *P* (*t*) ∝ *t^β^* exp (−*t/τ*). The fitted power-law exponent, *β* = −1.5, is consistent with the first-passage-time distribution expected for one-dimensional Brownian diffusion^82^ and agrees with previous findings ^39^. In the present slab geometry, this behaviour can arise because exchange between condensate regions is primarily governed by diffusion along the direction normal to the slab, while the exponential cutoff reflects the finite characteristic residence time.

To quantify differences in protein diffusivity between the two regions, we compute the PDF of the squared displacements of protein COM, Δ*r*^2^, for a lag time of *δt* = 20 ns, chosen such that proteins remained within the same region throughout the entire time interval *δt*. As expected, protein motion is more restricted in the bulk region than at the interface, as shown in Fig. 6b for WT and other sequences in Fig. S15.

Since some proteins remain in the condensate interior (cond. interior), whereas others are mobile and transition between the condensate interface and interior (cond. mobile), we compute the MSD over a larger time window of the protein COM for these two populations and compare them with the dilute-regime results in Fig. 6c–f. As expected, protein displacement in the dense phase is strongly suppressed compared with infinite dilution (see Fig. 6c).

In the dilute regime, MSD increases linearly with time, corresponding to a diffusive exponent close to unity (Fig. 6d). We compute the translational diffusion coefficient *D* from the Stokes–Einstein (SE) equation^83^ and apply the Yeh–Hummer correction^84,85^ accounting for finite-size effects arising from the finite simulation box (see Methods for the details). We obtain *D* values for WT, −7Y+7F, and −12F+12Y in a narrow range between 10.0 and 10.6

Å^2^ /ns, with −12F+12Y displaying the fastest diffusion. The values of the diffusion coefficients for WT and −7Y+7F differ by less than 0.1 A /ns, which is within statistical uncertainty of our estimations. For the +12E sequence, we find a slightly lower diffusion coefficient of 7.1 A Å^2^/ns, as this sequence was simulated at 277 K. For comparison, we also compute the theoretical expectation of the diffusion coefficient predicted by the SE relation, as shown in Fig. S16 in the SI. The theoretical values are of the same order as the ones extracted from simulations, however are larger by 20 % - 30 %. Such deviations are not unexpected, given the approximation of an IDP as a spherical particle in the SE description. When compared with experiments, these diffusion coefficients are of the same order as values expected for disordered proteins in aqueous solution. For example, the highly charged IDP ProT*α* with *R*_g_ ≈ 4.4 nm, has an experimental translational diffusion coefficient of approximately 8.5 Å^2^*/*ns at room temperature, with a comparable value obtained from atomistic simulations ^24^.

In contrast, the diffusion of a chain in protein condensates displays stronger sequence-specificity (see Fig. 6e,f). In a protein-rich environment, the chain motion is subdiffusive with the exponent *α* ≈ 0.75 for both interior and mobile regions, highlighting contact formation between chains that hinders their motion. When comparing with the results of Yamamoto and coworkers^39^, the obtained exponent is in agreement with subdiffusive motion of MDPs at short timescales, while for IDP condensates the authors recover the exponent *α* ≈ 0.85. This discrepancy might originate from the implicit treatment of the solvent in the CG model used in Ref. 39, which could lead to enhanced chain mobility.

We estimate an effective long-time diffusion coefficient, *D*_eff_, beyond the subdiffusive time window directly sampled in our multi-chain simulations. Assuming that the subdiffusive mean-square displacement, ⟨Δ*r*^2^(*t*)⟩ = *A*_app_*t^α^*, crosses over to normal diffusion once a protein has displaced over a distance comparable to its size, we define the crossover time as 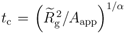. The extracted crossover times are of the same order as the trajectory length available for analysis, rationalizing why subdiffusive motion of the protein COM is observed over the timescales accessible in our simulations. We then estimate the corresponding long-time diffusion coefficient as 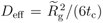. Here, *R̃*_g_ is the median radius of gyration of proteins assigned to the corresponding condensate region, and *A*_app_, *α* are the fitting parameters to the curves in Fig. 6e,f. The estimated diffusion coefficient in the condensate interior, 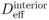, ranges from 0.004 to 0.046 Å^2^*/*ns for the +12E and −12F+12Y variants, respectively. Proteins in the mobile region diffuse faster, with 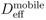 ranging from 0.016 to 0.124 Å^2^ */*ns for the same variants. These estimates fall within the broad range of experimentally and computationally reported protein mobilities in dense protein environments. For example, ProT*α* diffusion in the dense phase was reported to be approximately 0.27 Å^2^*/*ns^24^, which is comparable to the upper end of our mobile-region estimates but larger than most of our interior values. Di Bari *et al.*^86^ reported center-of-mass diffusion coefficients of approximately 0.1–0. 5 Å^2^*/*ns for unfolded proteins in a highly concentrated environment designed to mimic the *E. coli* cytoplasm after thermal stress. These values overlap primarily with the faster end of our estimates, although that study probes a crowded cytoplasmic environment rather than a phase-separated condensate and evaluates diffusion over a substantially shorter time regime. By contrast, single-molecule measurements of guest IDPs in p53 condensates yielded diffusion coefficients of approximately 0.003–0. 007 Å^2^*/*ns^87^, close to the slowest interior diffusion estimated here. This comparison is nevertheless not direct, because the experiments probe guest proteins diffusing through a p53 scaffold, whereas the A1-LCD molecules in our simulations constitute the condensate network itself.

Since Phe is more prone to contact formation than Tyr in the M3-IDP-C model, protein mobility increases as Phe residues are substituted with Tyr. Accordingly, the effective diffusion coefficient is the largest for the sequence with no Phe residues, −12F+12Y, intermediate for WT, which contains 12 Phe and 7 Tyr residues, and smallest for the Phe-rich sequence, −7Y+7F, which contains 19 Phe residues, in both condensate regions (Fig. 6e,f). In the inset of Fig. 6f, we further find a linear relationship between protein size in solution, extracted from the single-chain simulations shown in Fig. 5b, and the effective center-of-mass diffusion coefficient of proteins located in the condensate interior. For this comparison, we used the three sequences simulated under the same conditions that share nearly identical physicochemical properties: WT, −12F+12Y, and −7Y+7F. Consistent with the hierarchy of interaction strengths in M3-IDP-C, −7Y+7F adopts the most compact conformations in solution, forms the longest-lived residue contacts in condensates, and displays the slowest apparent diffusion. This trend supports the idea that the same interactions that compact isolated chains in solution can also promote longer-lived interchain contacts in the dense phase, thereby linking single-chain conformations to condensate dynamics ^66^. Similar relationships between single-chain and multichain properties have been reported for other IDP systems. For example, Mittal and coworkers investigated the material properties of model and natural highly charged polyampholytic IDPs^40^. Using HPS and Martini 3 models to describe biomacromolecules, they showed that protein size in solution correlates with condensate properties such as residue diffusion coefficients, surface tension, and condensate viscosity. This relationship was rationalized by the fact that stronger and longer-lived intrachain contacts lead to more compact protein conformations in solution and, in the dense phase, manifest as longer-lived interchain contacts. Similar conclusions linking interchain contact lifetimes and the material properties of condensates formed by charged IDPs were reported by Galvanetto et al. using experiments and all-atom simulations^24,26^.

Taken together, the estimated diffusion coefficients of A1-LCD in both dilute solution and condensates fall within the broad range reported for intrinsically disordered proteins. Although coarse-grained models generally accelerate molecular motion relative to experiment, the present Martini model reproduces the correct order of magnitude of protein diffusion.

## Discussion

Biomolecular condensates organize proteins and nucleic acids into dynamic membraneless compartments, often through transient multivalent interactions involving intrinsically disordered proteins or regions^1–3^. Because condensate formation and dissolution are tightly coupled to cellular function and disease, understanding the physicochemical interactions that control their structure, dynamics, and solvent environment remains essential^4,9,10^.

Computational models provide a direct route to connect sequence-dependent interactions of intrinsically disordered proteins with the emergent properties of biomolecular condensates. In this context, explicit-solvent coarse-grained models occupy a unique position by retaining an explicit representation of solvent and ions while remaining computationally efficient enough to access the length and time scales relevant to multichain condensates. Developing such models, however, requires achieving a transferable balance between protein–protein, protein–solvent, and ion-mediated interactions that simultaneously reproduces the behaviour of disordered proteins in dilute solution and in the condensed phase. Here, we assessed the performance of the Martini3-IDP model ^50^ and investigated whether improving its description of single-chain conformations translates into a more accurate description of biomolecular condensates. Using the extensively characterized A1-LCD family as a benchmark^30,41,60,61,64–66^, we identify which condensate properties are improved by refining dilute-state conformations and which aspects remain limited by the current treatment of solvent- and ion-mediated interactions.

We first examined whether improving the description of dilute-state conformations leads to a more transferable explicit-solvent coarse-grained model for biomolecular condensates. Consistent with the established relationship between single-chain dimensions and phase behaviour^16,56,57^, the Martini3-IDP model systematically underestimated the dimensions of six A1-LCD variants. Minimal, physically motivated refinements to protein–water and glycine interactions yielded excellent agreement with experimental radii of gyration and substantially improved transferability across a benchmark of 20 intrinsically disordered proteins. We termed the improved model Martini3-IDP-C, reflecting its intended application to biomolecular condensates.

Having established satisfactory performance at the single-chain level, we then asked to what extent this improved description of proteins in dilute solution translates into a more realistic representation of condensed-phase properties, using the well-characterized A1-LCD-family biomolecular condensates to examine how aromatic residues and sequence charge shape condensate properties. The resulting condensates remained highly hydrated and exhibited protein densities that were substantially closer to experiment than those predicted by the Martini3-IDP model. At the same time, no stable dilute phase was observed, indicating that the balance between protein–protein, protein–water, and ion-mediated interactions remains shifted toward excessive condensation. Thus, while improving single-chain conformations clearly enhances the description of biomolecular condensates, it is not by itself sufficient to quantitatively reproduce phase equilibria^34^. Nevertheless, the simulations successfully captured several key physicochemical properties of A1-LCD condensates, including the partitioning of both cations and anions into the dense phase, in semi-quantitative agreement with recent experiments^65^, and predicted key interchain contacts responsible for condensate formation.

Proteins within the condensates adopted coil-like conformations and were more expanded than in dilute solution. This observation is consistent with the polymer-physics view of biomolecular condensates as semidilute polymer solutions ^26^. Indeed, the densities of the simulated protein-rich phases are above the overlap concentrations, *c^∗^* = 3*M*_W_*/*4*N*_A_*π*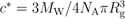, at which protein chains begin to interpenetrate^79^. For the IDPs studied here, with the molecular weight of *M*_W_ ≈ 13, 000g/mol and *R*_g_ ≈ 2.6 nm in the dilute regime, *c^∗^* falls in the range of approximately 300 mg/mL, while the dense phase remains substantially hydrated.

By contrast, the more compact conformations observed in solution likely reflect the poor effective solvent quality that promotes phase separation and condensate formation. Similar conclusions have been drawn from simulations using other coarse-grained models^34,38^, including the most recent version of the Martini3-IDP model^51,52^, suggesting that this behaviour is not specific to the Martini representation.

The Martini3-IDP-C model recapitulates key residue-level contact patterns associated with phase separation in the A1-LCD family, including contacts between aromatic residues and between aromatic and positively charged residues, in agreement with experimental observations^60,61^. Owing to the relatively fine resolution of the model, we further show that these contacts arise predominantly from side-chain–side-chain interactions. Although specific interactions such as *π*–*π* and cation–*π* interactions are not explicitly encoded in the Martini potential, these contact patterns are consistent with these interaction types.

Finally, we find that protein dynamics differ markedly between dilute solution and the condensed phase. In dilute solution, proteins undergo Brownian motion, with diffusion coefficients comparable to experimental values reported for proteins of similar size. By contrast, on the timescales accessible in our simulations, protein motion is subdiffusive both within the condensate interior and at the interface, consistent with the short-time dynamics observed in recent large-scale coarse-grained simulations of biomolecular condensates^39^. We attribute this behaviour to the formation of predominantly short-lived intermolecular contacts that transiently hinder protein mobility. Furthermore, we identify a linear correlation between the single-chain dimensions in dilute solution and protein diffusion within condensates, linking dilute-state conformations to the material properties of the condensed phase, in agreement with previous studies^40^. The resulting effective diffusion coefficients, estimated at the crossover time when a protein diffuses a distance comparable to its own size, are of the same order of magnitude as those reported in all-atom simulations and experimental measurements of intrinsically disordered proteins^24,86,87^.

Overall, our results demonstrate that Martini3-IDP-C can capture several key physicochemical properties of A1-LCD condensates in a semi-quantitative manner, including hydration, conformational behaviour, residue-level contact patterns, and protein dynamics. At the same time, our study highlights remaining limitations. In particular, the weak temperature dependence of standard Martini interactions limits the construction of phase diagrams over experimentally relevant temperature ranges^88^, while the treatment of solvent and ions may require further refinement to improve the balance between dilute- and dense-phase properties and ion partitioning. Addressing these challenges will be an important direction for future work and should further extend the applicability of the Martini3-IDP-C model to IDP phase separation and biomolecular condensates. In addition, the chemically detailed representation of Martini makes it well-suited for multiscale workflows in which equilibrated phase-separated configurations are backmapped to atomistic resolution, enabling detailed analysis of atomic-level interaction patterns while retaining access to larger-scale condensate organization^23,27^.

## Methods

### Simulation Details

We performed molecular dynamics (MD) simulations using the Martini3-IDP-C (M3-IDP-C) model developed in this work and the GROMACS simulation package^89,90^ (version 2023.5). M3-IDP-C is based on the recent M3-IDP model^50^, and includes two modifications: (*i*) strengthening of backbone (BB) bead–solvent interactions through the addition of a virtual site (VS) with *ε*_VS_*_−_*_W_ = 0.11 kJ/mol; (*ii*) reassignment of the glycine backbone bead from SP1 to SP1h to enhance glycine self-interactions. We also adjusted the equilibrium backbone angle (BBB) from 137*^◦^* to 130*^◦^* to improve model stability across different simulated temperatures.

To set up the single-chain simulations, we first used the AlphaFold server^91^ to generate all-atom starting configurations from the protein sequences. The resulting structures were then converted to coarse-grained representations and corresponding topology files using Martinize2^92^. Following the previous development of the model^50^, the secondary structure was set to *D* for all residues. Each protein was initially placed in a dodecahedral box and subjected to a short energy minimization. The system was then solvated, and sodium and chloride ions were added at concentrations corresponding to the experimental conditions, followed by energy minimization of the full system.

Each system was equilibrated by a short NpT simulation using the Berendsen thermostat and barostat to maintain the target temperature and pressure of 1 bar^93^. For production simulations, temperature was controlled with the stochastic velocity-rescaling thermostat^94^ applied separately to protein and non-protein groups, using a coupling constant of 1.0 ps. Pressure coupling was performed with the Parrinello–Rahman barostat^95^ in the isotropic ensemble, using a coupling constant of 12 ps and a target pressure of 1 bar. Production simulation times ranged from 10 to 20 *µ*s, depending on protein chain length. Sequence names, chain lengths, simulation conditions (temperature, salt concentration, and box size), and reference and simulated radii of gyration obtained with M3-IDP and M3-IDP-C are reported in Table S1 of the Supporting Information.

For the slab simulations, we first randomly inserted 200 copies of each protein into a rectangular simulation box with initial dimensions of 17 × 17 × 120 nm^3^, following previous recommendations for A1-LCD condensate simulations to reduce finite-size effects ^41^. The inserted protein conformations were selected from the corresponding single-chain trajectories and had radii of gyration close to the average value obtained under dilute conditions. To avoid the formation of multiple nucleation sites during condensate formation, we prepared a single dense phase by applying flat-bottomed position restraints to protein backbone beads along the *z*-axis, thereby concentrating the chains in the middle of the simulation box at the beginning of the simulation. We verified that this procedure did not produce overly compact dense phases by comparing it with a control simulation in which the slab formed from randomly dispersed chains without applying the flat-bottomed potential. As shown in Fig. S3, the mass and charge density profiles are indistinguishable between the two simulations. Computational details and additional discussion of this procedure are provided in the Supplementary Information.

The remaining system setup followed the single-chain protocol, with the exception that semi-isotropic pressure coupling was applied only along the slab normal, i.e., the *z*-axis, while the lateral box dimensions were kept fixed. Thus, the simulations were performed in the *NTAp_z_* ensemble, where *A* = *L_x_L_y_*. Production simulations were run for at least 30 *µ*s. We used standard MD settings for the Martini force field^46^. The integration time step was 20 fs. Van der Waals interactions were calculated using a potential-shift scheme with a cutoff of 1.1 nm, and Coulomb interactions were treated using a reaction-field method with a relative dielectric constant of 15 and a cutoff of 1.1 nm.

### Trajectory analysis

We performed trajectory analysis using GROMACS analysis tools and the MDAnalysis library (version 2.9.0)^96,97^. Mass and charge density profiles were computed using the *gmx density* command. Because bead masses in Martini 3 do not exactly correspond to the atomic masses of the mapped atoms, the resulting protein mass densities were rescaled by the ratio between the protein molecular weight and its corresponding Martini 3 bead-based molecular weight. All other analyses were performed using MDAnalysis.

#### Sequence Descriptors

We used the following descriptors of a protein sequence composed of *N* amino acids to characterize its physicochemical properties: (1) the fraction of aromatic residues, 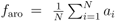, where *a_i_* = 1 for phenylalanine (F), tyrosine (Y), and tryptophan (W); (2) the net charge per residue, 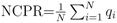, where *q_i_* is the charge per residue; (3) the sequence charge decoration^72^, 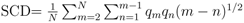, where *q_m_* and *q_n_*are charges at position *m* and *n*, respectively; (4) the average residue stickiness 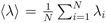, where *λ_i_* is the recently developed stickiness scale for disordered protein^73^, which was derived to reproduce the experimental data of a large set of disordered proteins. Here, we used the *λ_i_* values that were derived assuming the mean size of a residue is *σ*=0.56 nm.

#### Contact map calculations

We adopted a bead-size- and bead-type-dependent definition of interchain contacts. Specifically, two proteins were considered to be in contact if the minimum distance between any pair of beads *i* and *j*, belonging to residues on different protein chains, was smaller than 1.2*σ_ij_*, where *σ_ij_* is the Lennard-Jones radius representing the excluded volume of the corresponding bead pair. The factor of 1.2 was used to identify significant bead-pair interactions^41^. This choice was motivated by the fine-grained nature of the Martini force field, in which beads within a residue can have three different sizes: 0.47 nm (regular), 0.41 nm (small), and 0.34 nm (tiny). Backbone beads are typically represented by regular or small beads, whereas side chains are represented by small or tiny beads. We further decomposed interchain contacts into backbone–backbone, backbone–side-chain, and side-chain–side-chain contributions, which is not possible in one-bead-per-residue coarse-grained models.

The relative interchain contact probability was normalized by (*i*) the number of trajectory snapshots, (*ii*) the number of possible protein pairs, *N*_ch_(*N*_ch_ − 1)*/*2, and (*iii*) the mean residue–residue interchain contact probability, to highlight the relative contributions of backbone and side-chain bead pairs to residue–residue contact maps. Contact lifetimes were computed by tracking the continuous persistence of each interchain residue–residue contact along the trajectory. The last 10 *µ*s of each trajectory were used for this analysis.

To compute a composition-normalized enrichment of significant residue-type contact pairs *E_ij_*, we first define a contact pair as *significant* if its contact probability is greater than three times the mean residue-residue contact value. Next, we count the occurrence of this contact pair, ignoring residue indexes. Finally, we compute the enrichment of a contact formed by *i* − *j* pair as 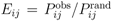, where 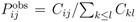 is the observed fraction of significant contacts of *i* − *j* type, and the expected probability assuming random pairing of an *i* − *j* contact pair is defined as 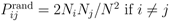 and 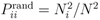 otherwise, where *N_i_* is the number of residues of type *i* in the sequence, *N* is the chain length, and the factor 2 accounts for interchangeability of *i* − *j* pair.

#### Mean-squared displacement calculations

For a single protein in solution, we computed the mean-squared displacement (MSD) of its center of mass (COM). Before calculating COM trajectories, we ensured that the protein was not split across periodic boundaries in each trajectory frame. For proteins partitioned into the condensate, we computed the MSD of the COM of each protein in the reference frame of the condensate. Specifically, before calculating displacements, we subtracted the translational motion of the protein-rich phase. Apparent diffusion coefficients were extracted by fitting the MSDs over the 2–50 ns time window.

For single chain simulations, we estimate the translational diffusion coefficient from the Einstein relation, ⟨Δ*r*^2^(*t*)⟩ = 6*D*_PBC_*t*, where *D*_PBC_ is the diffusion coefficient obtained from simulations under periodic boundary conditions (PBC). The diffusion coefficients were extracted by fitting the MSDs over the 2–50 ns time window. To account for hydrodynamic finite-size effects arising from the finite simulation box, we applied the Yeh–Hummer correction^84,85^, 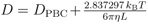, where *L* is the cubic box length and *η* is the shear viscosity of Martini water at the corresponding temperature. The viscosity is determined independently from equilibrium simulations using the Green–Kubo formalism^98^. All single-chain diffusion simulations were performed in the NVT ensemble using the mean box dimensions extracted from previous NPT simulations to avoid possible systematic errors in diffusion coefficients introduced by box-volume fluctuations in NPT simulations ^99^.

## Data availability

The data that support the findings of this study are available from the corresponding author upon reasonable request.

## Acknowledgments

T.I.M. acknowledges support by the French National Research Agency (Agence Nationale de la Recherche, ANR) through the project DiPCaM, project number ANR-25-CE06-0062-01. L.B.-A., P.C.T.S and T.I.M. acknowledge support from the French National Center for Scientific Research (CNRS). P.C.T.S also acknowledges funding through research collaboration agreements with Sanofi. K.L.-L. acknowledges support by the Novo Nordisk Foundation via the PRISM (Protein Interactions and Stability in Medicine and Genomics) centre (NNF18OC0033950). This work was performed using HPC resources from GENCI–IDRIS, GPU-accelerated partitions of the Jean Zay supercomputer (Grant 2021-2026 - A0100712464). The authors also acknowledge the support of the Centre Blaise Pascal’s IT test platform at ENS de Lyon (Lyon, France) for the computer facilities. The platform operates the SIDUS solution developed by Emmanuel Quemener ^100^.

## Competing interests

K.L.-L. holds stock options in and is a consultant for Peptone. The remaining authors declare no competing interests.

