## Supporting Information for "Towards transferable explicit-solvent coarse-grained models for biomolecular condensates"

†*Laboratoire de Physique de l'École Normale Supérieure de Lyon (LPENSL), CNRS UMR 5672, École Normale Supérieure de Lyon, 69342 Lyon Cedex 07, France*

‡*Centre Blaise Pascal de Simulation et de Modélisation Numérique, École Normale Supérieure de Lyon, Lyon, France*

¶*Laboratoire de Biologie et Modélisation de la Cellule, CNRS UMR 5239, Inserm U1293, Université Claude Bernard Lyon 1, École Normale Supérieure de Lyon, Lyon, France*

§*Department of Biology, Structural Biology and NMR Laboratory, Linderstrøm-Lang Centre for Protein Science, University of Copenhagen, 2200 Copenhagen, Denmark*

*

### Single-chain simulations

Table 1: Single-chain simulation conditions and radii of gyration for the reference data and the M3-IDP and M3-IDP-C models. Sequences shorter than 140 residues were simulated for 10 $\mu$s, while larger sequences were simulated for 20 $\mu$s. Values of $R_g$ are reported as mean $\pm$ error for the reference data and mean $\pm$ SEM for simulations.

| Protein | $N_{res}$ | $T$ (K) | $c_{salt}$ (M) | $L_{box}$ (nm) | $R_g^{ref}$ (nm) | $R_g^{M3\text{-}IDP}$ (nm) | $R_g^{M3\text{-}IDP-C}$ (nm) | Reference |
|---|---|---|---|---|---|---|---|---|
| WT | 137 | 298 | 0.150 | 23.5 | $2.76 \pm 0.02$ | $2.24 \pm 0.04$ | $2.66 \pm 0.05$ | 1 |
| WT+NLS | 137 | 298 | 0.150 | 23.5 | $2.58 \pm 0.01$ | $2.11 \pm 0.03$ | $2.60 \pm 0.07$ | 1 |
| -10R+10K | 137 | 298 | 0.150 | 23.5 | $2.85 \pm 0.01$ | $2.12 \pm 0.03$ | $2.61 \pm 0.07$ | 1 |
| -6R+6K | 137 | 298 | 0.150 | 23.5 | $2.79 \pm 0.01$ | $2.18 \pm 0.02$ | $2.65 \pm 0.09$ | 1 |
| -12F+12Y | 137 | 298 | 0.150 | 23.5 | $2.60 \pm 0.02$ | $2.25 \pm 0.07$ | $2.63 \pm 0.04$ | 1 |
| -7Y+7F | 137 | 298 | 0.150 | 23.5 | $2.72 \pm 0.01$ | $2.14 \pm 0.04$ | $2.79 \pm 0.08$ | 1 |
| +12D | 137 | 298 | 0.150 | 23.5 | $2.80 \pm 0.01$ | $2.01 \pm 0.03$ | $2.31 \pm 0.06$ | 1 |
| +12E | 137 | 298 | 0.150 | 23.5 | $2.85 \pm 0.01$ | $2.34 \pm 0.08$ | $2.79 \pm 0.07$ | 1 |
| +7K12D | 137 | 298 | 0.150 | 23.5 | $2.92 \pm 0.01$ | $2.07 \pm 0.02$ | $2.45 \pm 0.06$ | 1 |
| K25 | 185 | 288 | 0.150 | 25.0 | $4.10 \pm 0.2$ | $2.56 \pm 0.04$ | $3.05 \pm 0.07$ | 2 |
| K44 | 283 | 288 | 0.150 | 32.0 | $5.20 \pm 0.2$ | $2.73 \pm 0.04$ | $3.20 \pm 0.05$ | 2 |
| PNt | 334 | 298 | 0.150 | 35.0 | $5.11 \pm 0.10$ | $3.31 \pm 0.11$ | $5.23 \pm 0.27$ | 3 |
| PNtS3 | 334 | 298 | 0.150 | 35.0 | $4.06 \pm 0.10$ | $3.30 \pm 0.11$ | $4.85 \pm 0.22$ | 3 |
| $\alpha$Syn140 | 140 | 293 | 0.200 | 23.5 | $3.55 \pm 0.10$ | $3.87 \pm 0.11$ | $4.31 \pm 0.10$ | 4 |
| N-CoRNID | 271 | 293 | 0.192 | 35.0 | $4.70 \pm 0.2$ | $2.78 \pm 0.05$ | $3.29 \pm 0.07$ | 5 |
| GLY150 | 150 | 290 | 0.000 | 27.0 | $1.47 \pm 0.05$ | $2.99 \pm 0.03$ | $2.58 \pm 0.02$ | 6 |
| GLY200 | 200 | 290 | 0.000 | 27.0 | $1.60 \pm 0.05$ | $3.50 \pm 0.03$ | $2.89 \pm 0.03$ | 6 |
| ELP3 | 18 | 295 | 0.100 | 10.0 | $0.99 \pm 0.01$ | $1.18 \pm 0.00$ | $1.15 \pm 0.00$ | 7 |
| ELP45 | 228 | 295 | 0.100 | 30.0 | $2.17 \pm 0.10$ | $3.49 \pm 0.10$ | $4.20 \pm 0.10$ | 7 |
| $\beta$-casein | 209 | 300 | 0.000 | 25.0 | $2.30 \pm 0.10$ | $2.80 \pm 0.08$ | $4.58 \pm 0.14$ | 8 |

#### Relaxation timescales of a single protein

To ensure that we simulated single-chain systems long enough, we compute the ratio between the simulation time, $t_{\mathrm{sim}}$, and the characteristic timescales of a single chain. In particular, we quantify the orientation relaxation of a protein by computing the autocorrelation function (ACF) of its end-to-end vector $\mathbf{R}_{\mathrm{ee}}$. To describe size fluctuations of a chain, we compute the ACF of the square radius of gyration $R_{\mathrm{g}}^2$. For both ACFs, we estimate the integrated autocorrelation time as $\tau_i = 1 + 2\sum C_i$, where $i$ is either $\mathbf{R}_{\mathrm{ee}}$ or $R_{\mathrm{g}}^2$, and we sum until the first point where the value of $C_i$ becomes negative. In conclusion, we simulate single-chain systems for at least twenty times longer than their longest relaxation timescale.

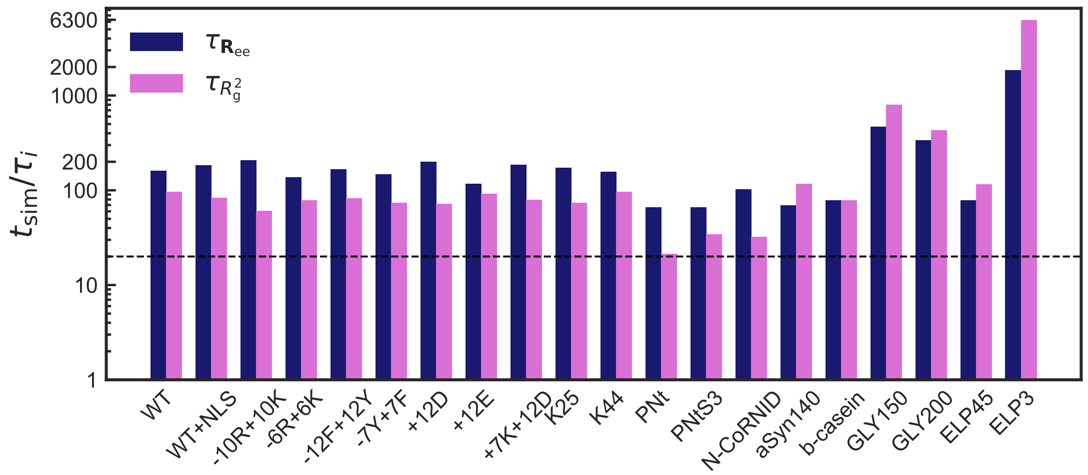


Figure 1: Ratio between the simulation time, $t_{\mathrm{sim}}$, and the single chain relaxation timescale computed from the ACFs of the end-to-end vector, $\tau_{\mathbf{R}_{\mathrm{ee}}}$, and the square radius of gyration, $\tau_{R_{\mathrm{g}}^2}$, shown in blue and pink, respectively. Dashed black line corresponds to $t_{\mathrm{sim}}/\tau_i$=20.

#### Preparation protocol for slab simulations

##### Seeding the dense phase

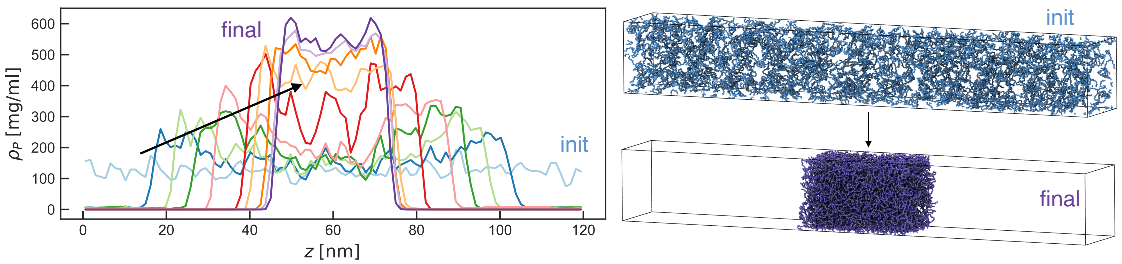


Figure 2: Temporal evolution of the WT protein mass density profile when the flat-bottomed (FB) position restraints are applied to backbone protein beads. Simulation snapshots at the beginning (init) and at the end of the simulations (final) are shown alongside. Only protein backbone beads are rendered for the sake of clarity.

Initially, we set up multi-chain simulations by randomly distributing proteins (RAN) within the simulation box. This approach generally resulted in the formation of multiple protein aggregates that did not merge even after long simulations exceeding 30 $\mu$s. The only exception was the WT sequence simulated at 298 K. This outcome is not unexpected, as the Martini force field treats solvent explicitly, which can slow down the diffusion of proteins and their assemblies compared with implicit-solvent coarse-grained models.

This result motivated us to develop an alternative approach in which the dense phase was prepared by applying flat-bottomed (FB) position restraints to protein backbone (BB) beads along the $z$-axis, thereby concentrating the chains in the middle of the simulation box at the beginning of the simulation. During a 1-$\mu$s simulation, we progressively adjusted the FB position restraints by reducing the distance from the centreof the box beyond which the restraints were applied. Specifically, $r_0$ was set to 30, 20, 10, 1, and 0.1 nm, and the force constant $k_{\text{FB}}$ was set to 0.05 kJ/mol/nm$^2$, except for $r_0 = 0.1$ nm, where it was reduced to 0.005kJ/mol/nm$^2$. The temporal evolution of the protein density profile for WT is shown in Fig. 2, together with initial and final system snapshots illustrating the formation of the protein slab. To evaluate whether our approach of seeding the dense phase has an influence on the later state of a system, we compare the mean mass and charge density profiles for a

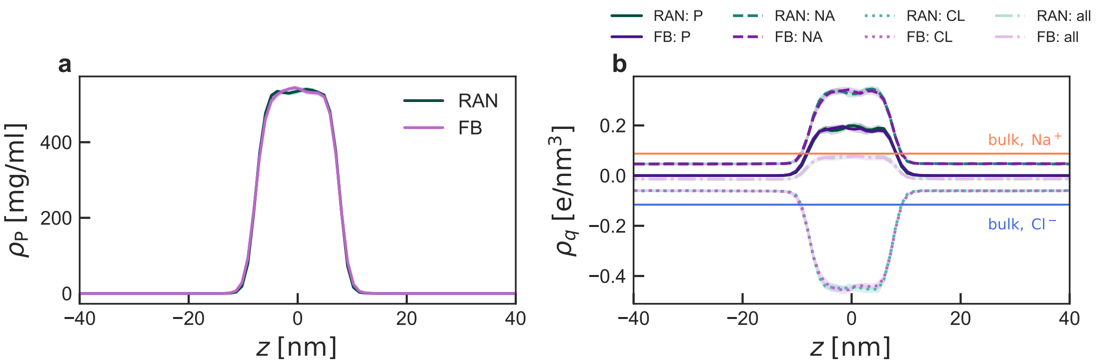


Figure 3: **a**: mean mass density profile of protein-rich phase when the starting configuration was prepared either by randomly (RAN) placing protein in a box or applying the flat-bottomed (FB) position restrains to protein backbone beads; **b**: mean charge density profile of protein (P), sodium cation ($Na^+$), chloride anion ($Cl^-$) beads, and the total charge density profiles along $z$-axis.

system composed of 110 proteins of WT sequence that formed a protein slab when starting from RAN and FB-mediated starting configurations shown in Fig. 3a and b. The curves for mass and charge densities are indistinguishable for both methods. Thus, we use the FB position restraints for preparing a protein-rich phase for all systems reported in the main text.

##### Detecting the dilute phase

We note that sampling the dilute phase is computationally challenging, as it corresponds to a small number of free protein chains dispersed in the simulation box for the systems studied here. We therefore first verified whether a dilute phase was observed in our simulations. Although the mean density profiles shown in Fig. 2 in the main text reach zero in regions far from the protein slab, rare events in which a protein leaves the dense phase could, in principle, occur and be averaged out in the density calculation. To exclude this possibility, we performed the following analysis. First, we defined the boundaries between the dilute and dense regions by fitting the average protein density profiles from Fig. 2 in the main text with a hyperbolic tangent function,

$$\rho_{\mathrm{P}}(z) = \frac{\rho_{\mathrm{dense}} + \rho_{\mathrm{dilute}}}{2} \pm \frac{\rho_{\mathrm{dense}} - \rho_{\mathrm{dilute}}}{2} \tanh\left(\frac{z - z_{\mathrm{DS}}}{d}\right), \quad (1)$$

where $\rho_{\mathrm{dilute}}$ and $\rho_{\mathrm{dense}}$ are the densities of the dilute and dense phases, respectively. The (+) sign was used to fit the density profile for $z > 0$, whereas the (-) sign was used for the $z < 0$ region. Here, $z_{\mathrm{DS}}$ and $d$ denote the position of the dividing surface and the interfacial thickness, respectively. We then defined the dilute region as $|z| > z_{\mathrm{DS}} + 2.5d$, where the prefactor 2.5 ensures that this region is sufficiently far from the dense phase[9]. Next, we compute the $z$-coordinates of the protein centers of mass (COM) over the equilibrated part of each trajectory and check whether any chain entered the dilute region. For all sequences studied here, we did not detect any protein chain in the dilute region.

**Seeding the dilute phase**

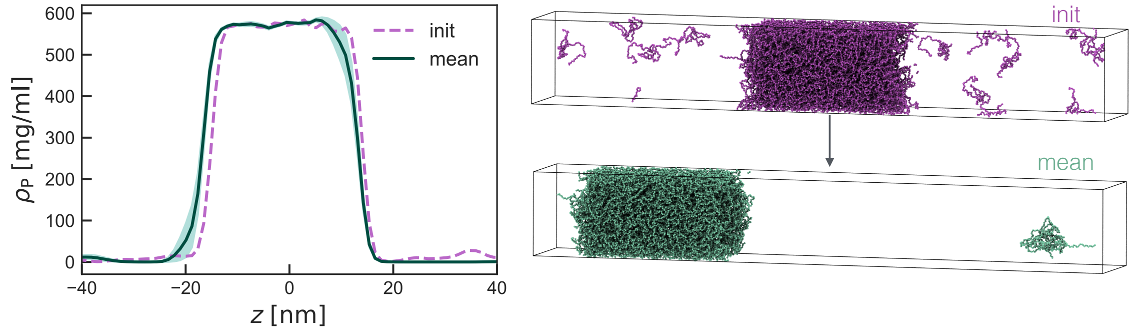


Figure 4: Mass density profile of the protein beads at the start (init) and averaged over the last ten microseconds (mean) along the $z$-axis, shown in purple and green, respectively. The corresponding system snapshots are shown on the right side of the figure using the same color scheme. Only protein backbone beads are rendered for the sake of clarity.

Because we did not observe a stable dilute phase over the course of long multichain simulations of the A1-LCD-family systems, exceeding 30 $\mu$s, we tested whether this behavior was due to limited simulation time or to overly strong protein-protein interactions, i.e., an intrinsic property of the force field. To this end, we set up a simulation in which the dilute phase was seeded directly. Specifically, we used the final configuration of the WT system composed of 200 chains forming a slab and added 10 additional protein chains by replacing solvent beads. Because each chain carries a net positive charge of $+8q_e$, we also added 80 additional $Cl^-$ ions by replacing solvent beads to maintain system electroneutrality. These system manipulations were performed using the GROMACS tool *gmx insert-molecules*. The resulting system was subjected to standard energy minimization, followed by a production simulation with semi-isotropic pressure coupling in the $Np_zT$ ensemble at $T = 293$ K. The simulation length was 20 $\mu$s.

In Fig. 4, we show the initial protein mass density profile along the $z$-axis together with the mean profile computed over the last 10 $\mu$s using 1-$\mu$s blocks. We also show simulation snapshots at time zero and after 20 $\mu$s. We observed that six out of ten free chains joined the dense phase, whereas the remaining chains formed a small aggregate in the dilute region. This aggregate is expected to eventually merge with the main dense phase to reduce the interfacial area with the solvent. Thus, the seeded dilute phase was unstable under these simulation conditions, supporting the absence of a stable dilute phase in the slab simulations reported in the main text.

### Mass density profiles in slab simulations of A1-LCD variants

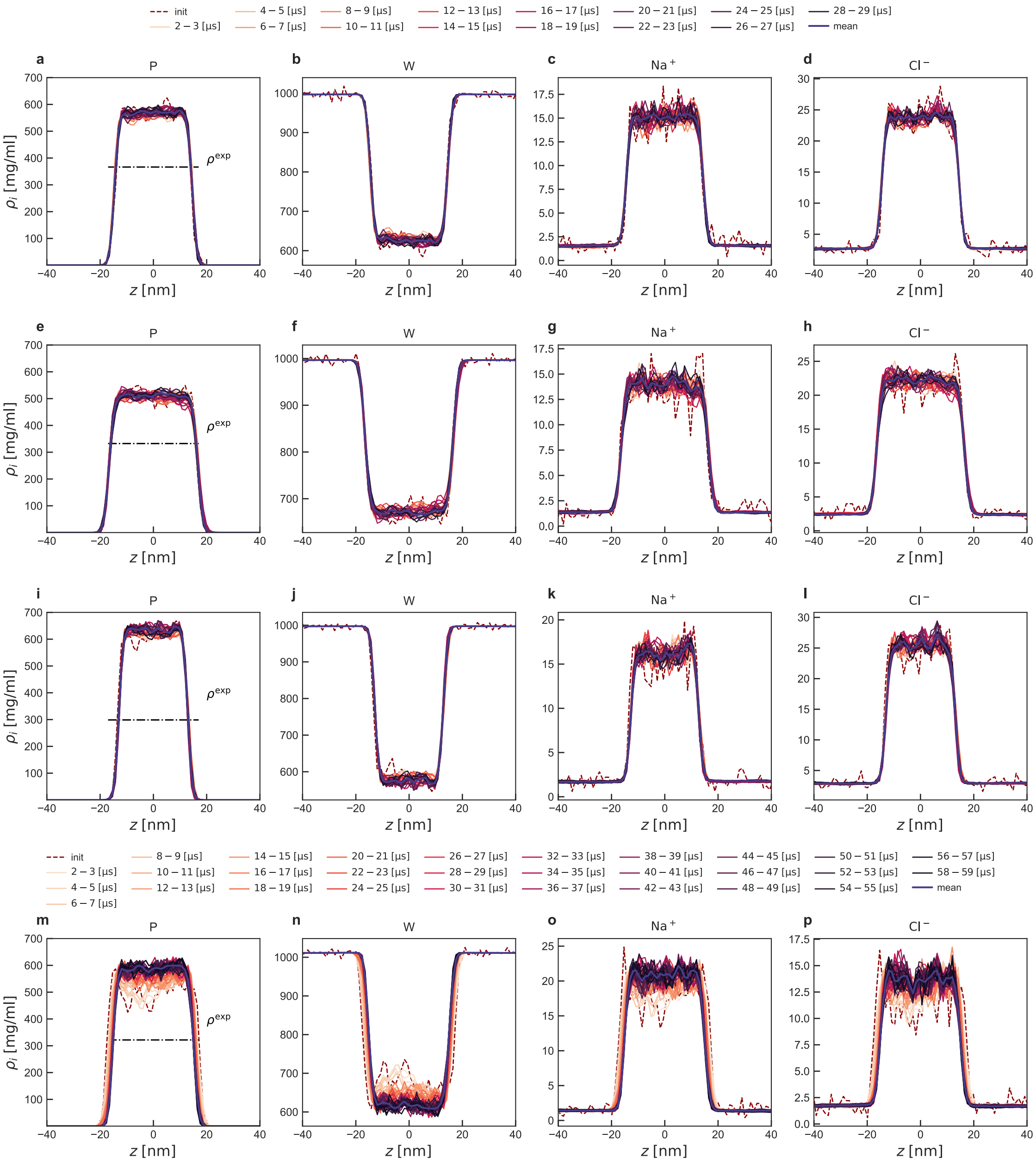


Figure 5: Temporal evolution of mass density profiles of protein (P), water (W), sodium cation ($Na^+$), and chloride anion ($Cl^-$) in WT (**a-d**), -12F+12Y (**e-h**), -7Y+7F (**i-l**), and +12E (**m-p**) in multi-chain systems. The initial and the mean density profiles are shown in dashed red and solid purple lines, respectively. Experimental protein densities in the dense phase, $\rho^{\mathrm{exp}}$, are shown as black dash-dotted lines.[1]

#### Comparison between protein-rich phase: M3-IDP *vs* M3-IDP-C

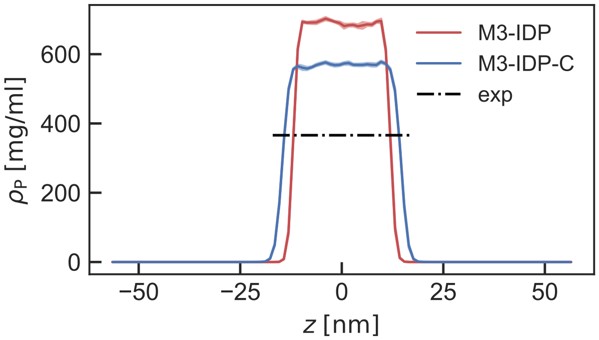


Figure 6: Mean mass density profile of the protein-rich phase for the WT sequence along the $z$-axis for M3-IDP[10] and M3-IDP-C model field shown in red and blue, respectively. Shaded areas represent the error bars, calculated as the standard error of the mean. The experimental density (exp) of the dense protein phase is shown as dash-dotted line.

#### Charge density profiles in slab simulations of A1-LCD variants

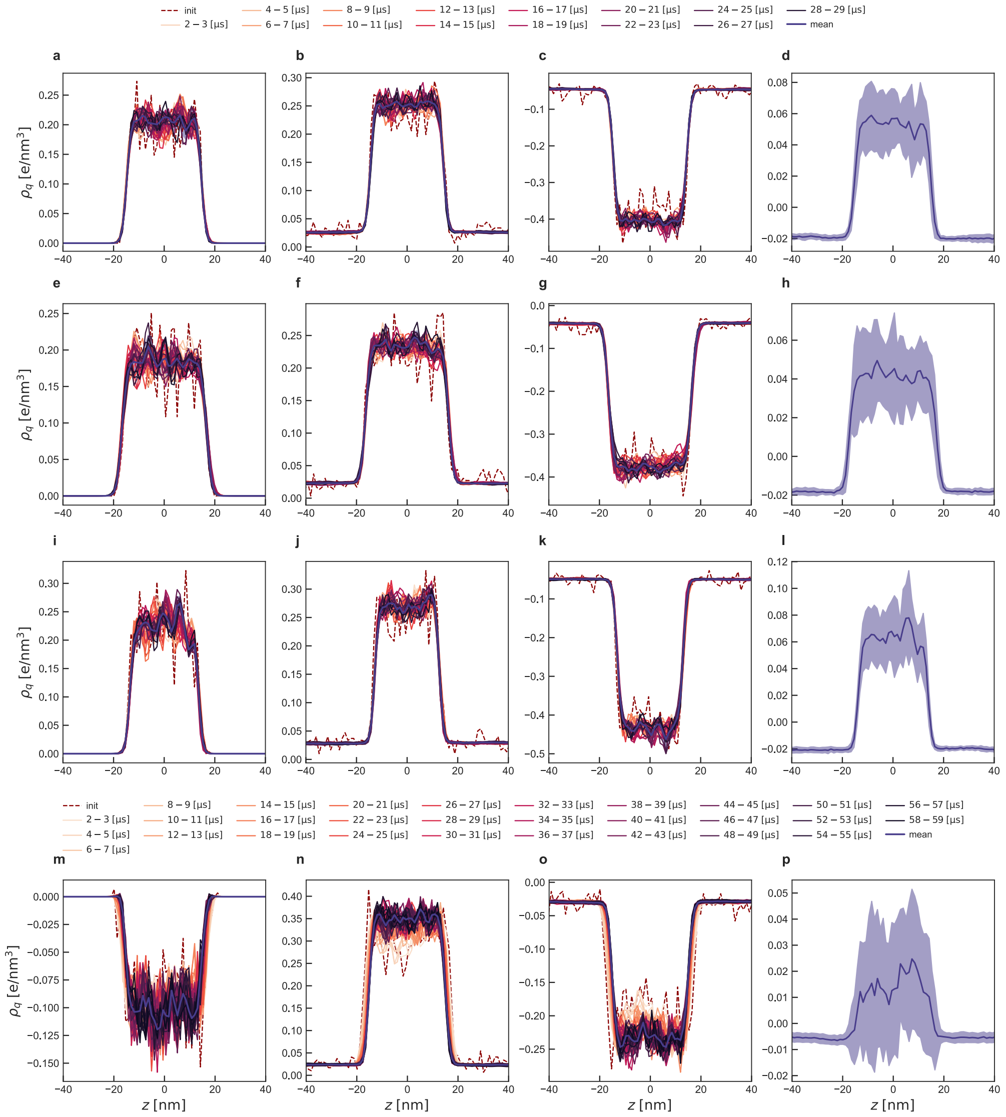

Figure 7: Temporal evolution of charge density profiles of protein (P), sodium cation ($Na^+$), chloride anion ($Cl^-$), and total charge density profiles in WT (**a-d**), -12F+12Y (**e-h**), -7Y+7F (**i-l**), and +12E (**m-p**) in multi-chain systems. The initial and the mean density profiles are shown in dashed red and solid purple lines, respectively.

#### Contact formation in A1-LCD sequences

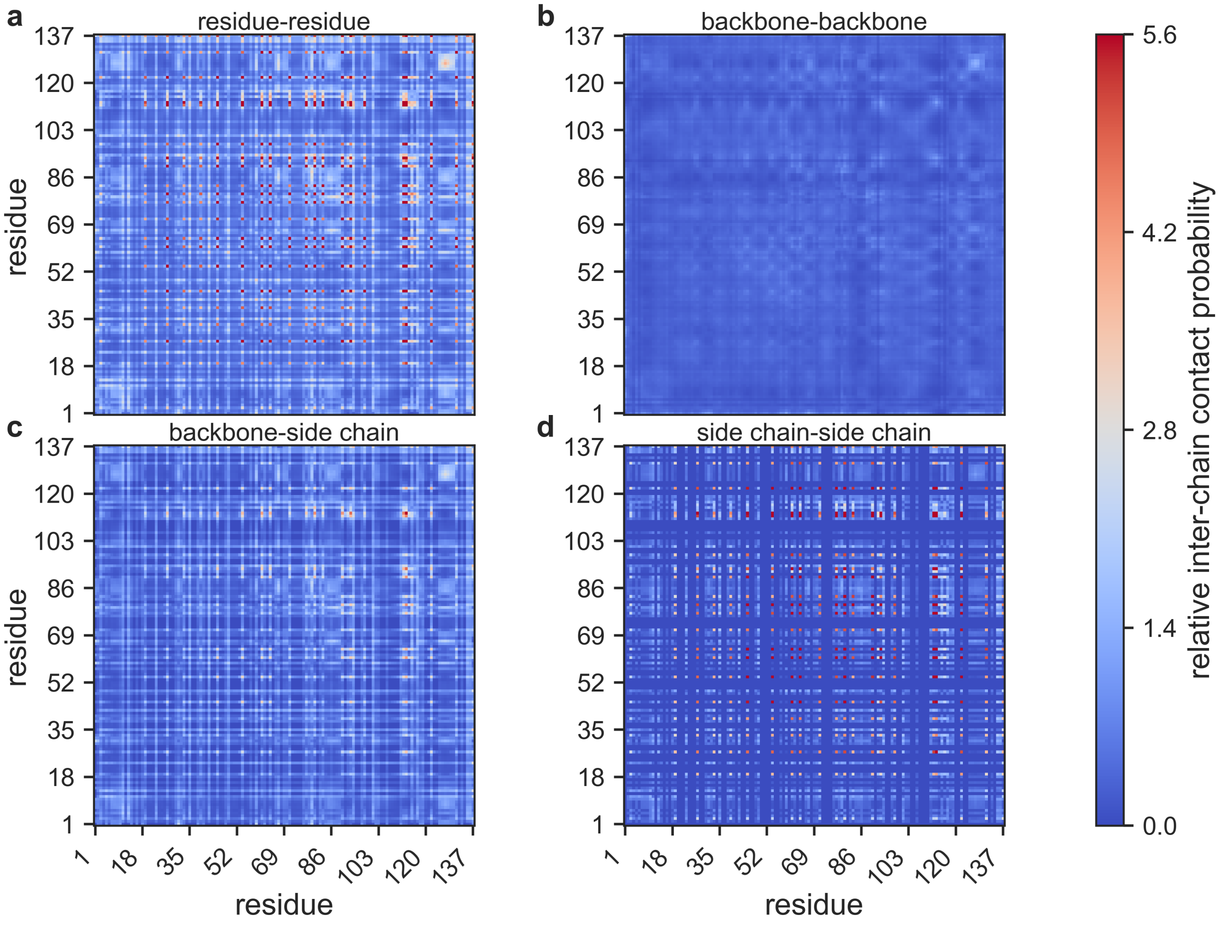


Figure 8: Inter-chain contact probability for -7Y +7F sequence **a-d**, using all, only backbone, backbone and side chain, and side chain beads. The absolute value is normalized by the number of possible chain pairs and the number of snapshots in a trajectory, and scaled by the mean value of the residue-residue contacts. The maximum value is capped at the 99th percentile of the relative residue-residue contacts.

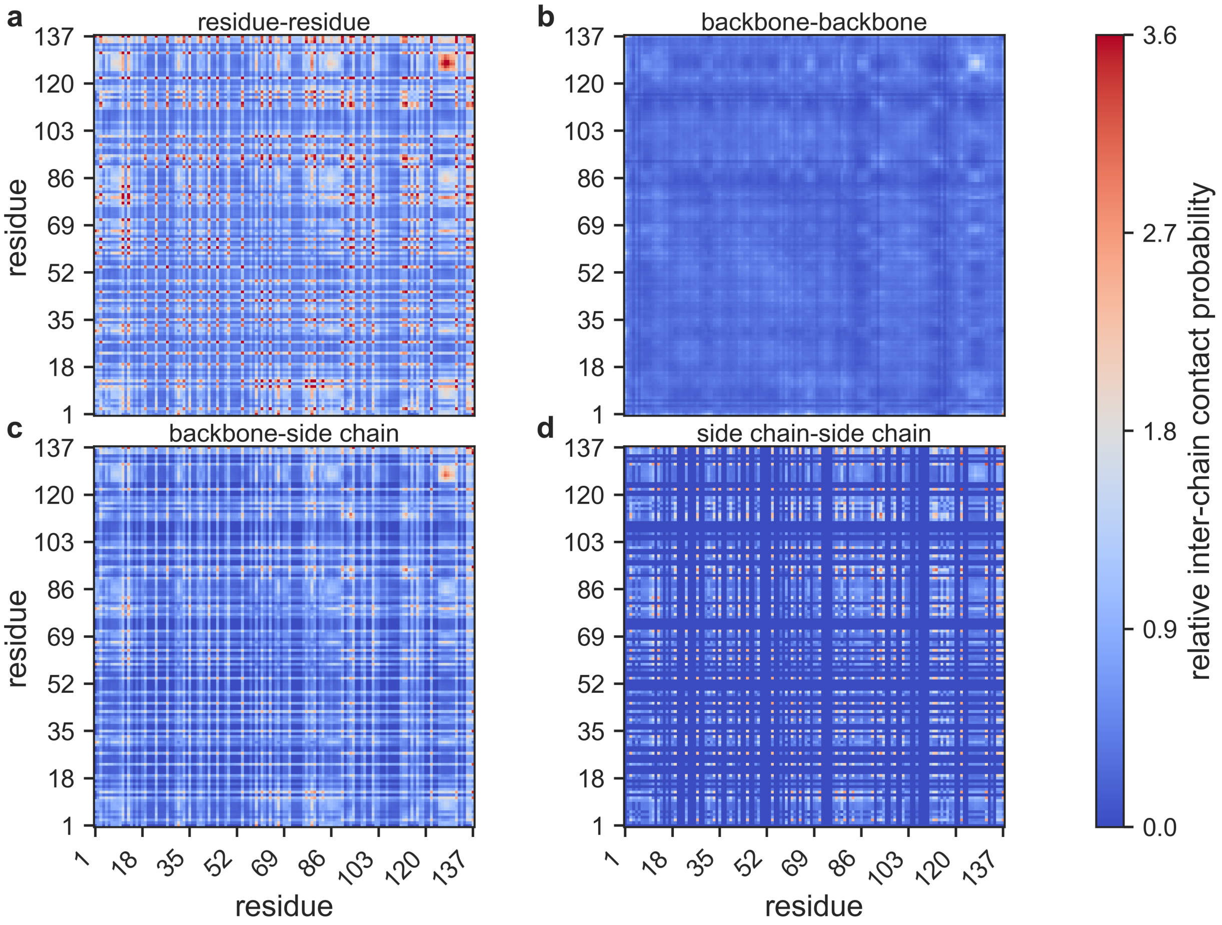


Figure 9: Inter-chain contact probability for -12F +12Y sequence **a-d**, using all, only backbone, backbone and side chain, and side chain beads. The absolute value is normalized by the number of possible chain pairs and the number of snapshots in a trajectory, and scaled by the mean value of the residue-residue contacts. The maximum value is capped at the 99th percentile of the relative residue-residue contacts.

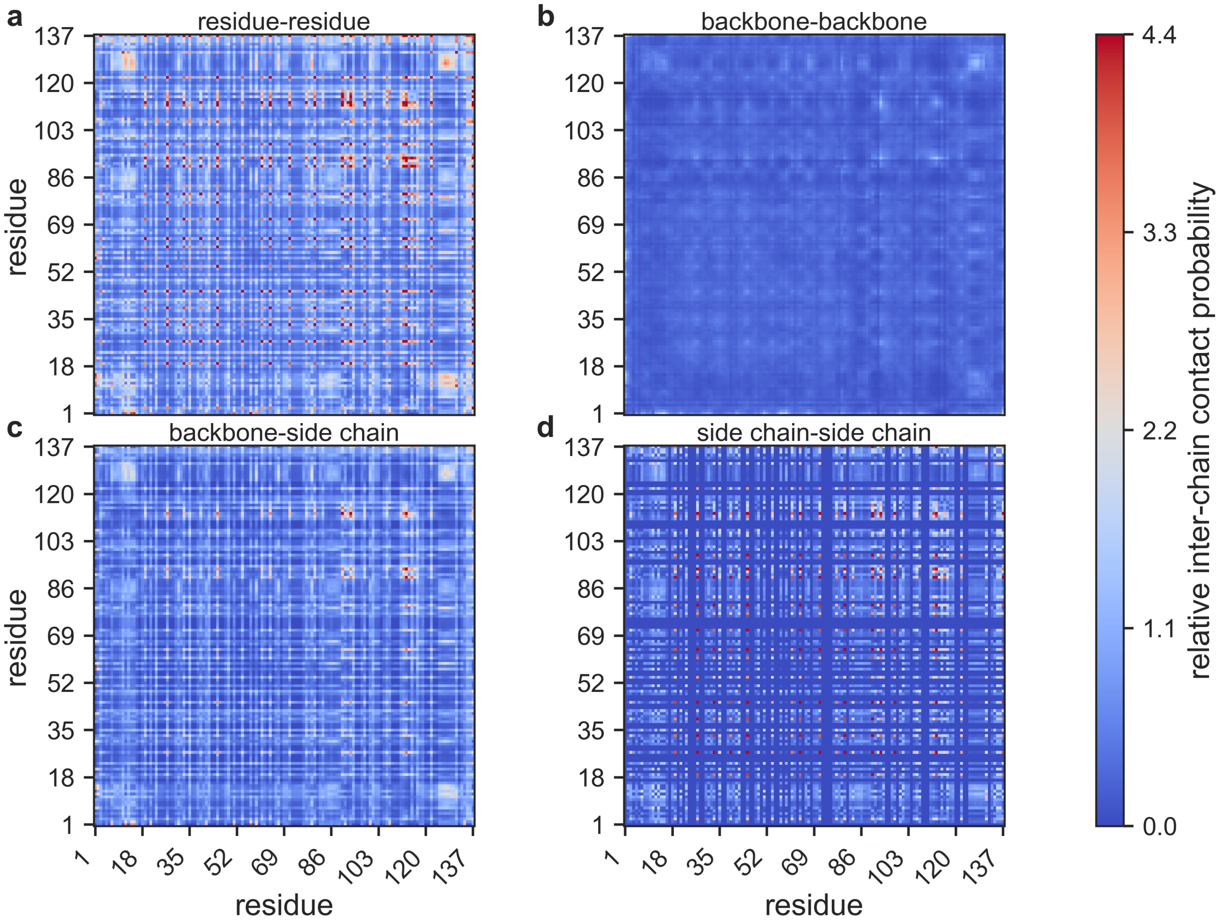


Figure 10: Inter-chain contact probability for +12E sequence **a**-**d**, using all, only backbone, backbone and side chain, and side chain beads. The absolute value is normalized by the number of possible chain pairs and the number of snapshots in a trajectory, and scaled by the mean value of the residue-residue contacts. The maximum value is capped at the 99th percentile of the relative residue-residue contacts.

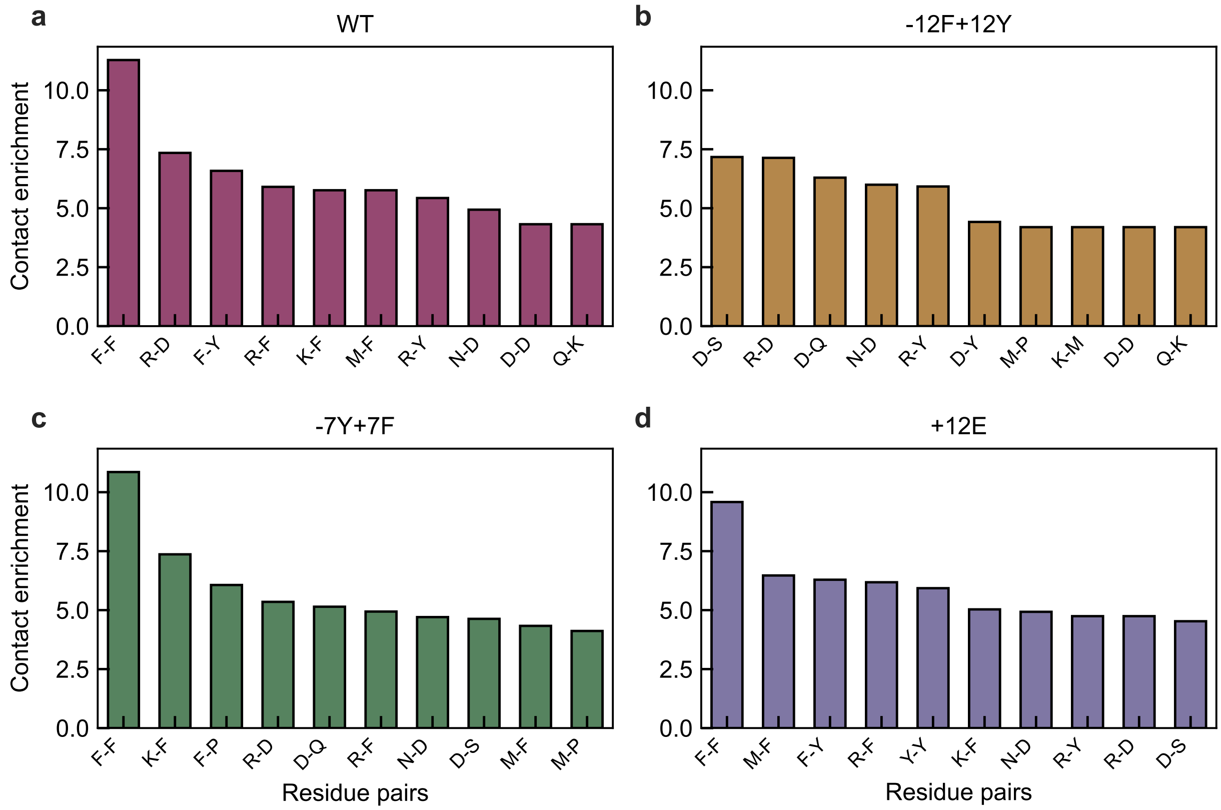


Figure 11: **a-d**, Enrichment of intra-chain contact pairs for WT,-12F+12Y, -7Y+7F, and +12E computed in single-chain simulations.

#### Protein size and conformations in solution and within condensates

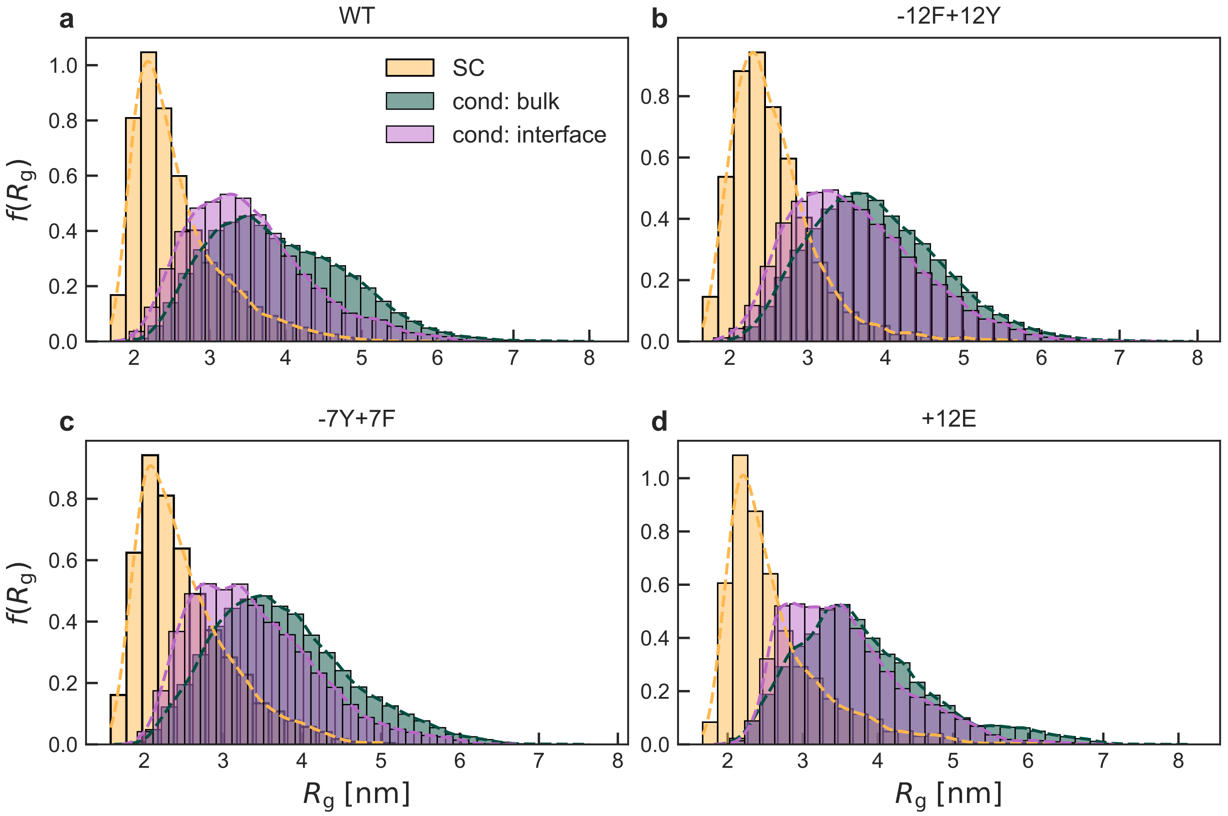


Figure 12: **a-d**, Probability density function (PDF) of the radius of gyration $f(R_\mathrm{g})$ for WT, -12F+12Y, -7Y+7F, and +12E in the dilute (single-chain, SC) limit, at the interface (cond. interface), and inside the protein-rich phase (cond. bulk).

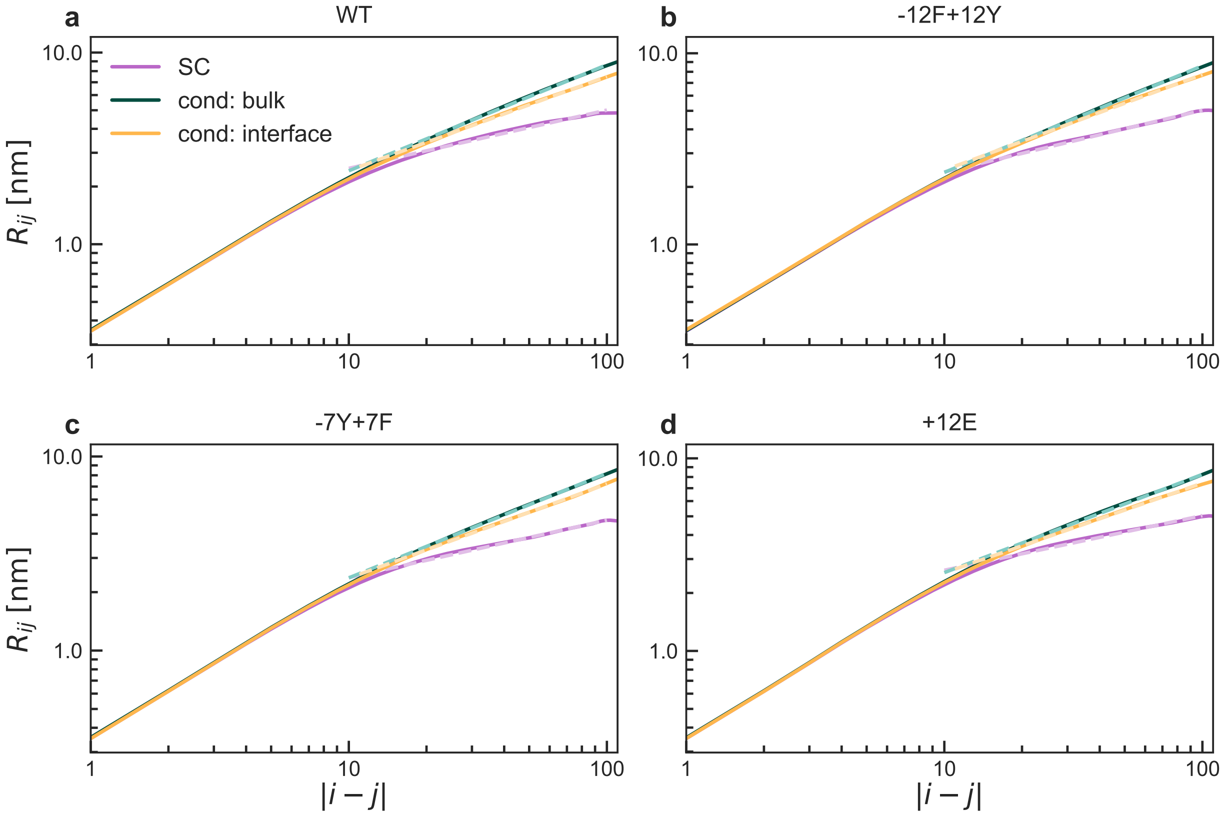


Figure 13: **a-d**, Root-mean-square distances $R_{ij}$ between residues $i$ and $j$ as a function of the distance along the sequence $|i-j|$ for WT, -12F+12Y, -7Y+7F, and +12E in the dilute (single-chain, SC) limit, at the interface (cond. interface), and inside the protein-rich phase (cond. bulk). Dashed curves correspond to the fits with the expression $R_{ij} = R_0|i-j|^{\nu^{\mathrm{app}}}$, where $\nu^{\mathrm{app}}$ is the apparent Flory exponent.

#### Protein dynamics in condensates

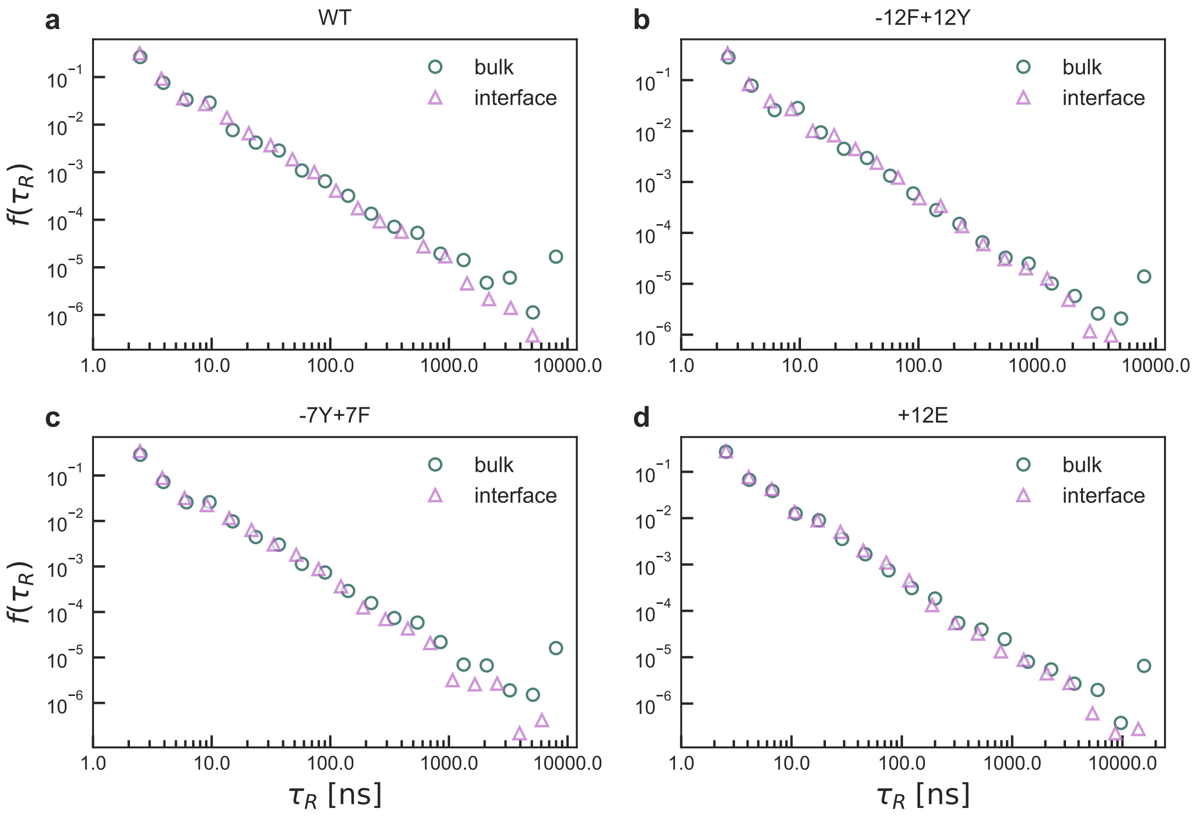


Figure 14: **a-d**, PDF of the residence time of A1-LCD proteins at the interface and inside the condensate for WT, -12F+12Y, -7Y+7F, and +12E.

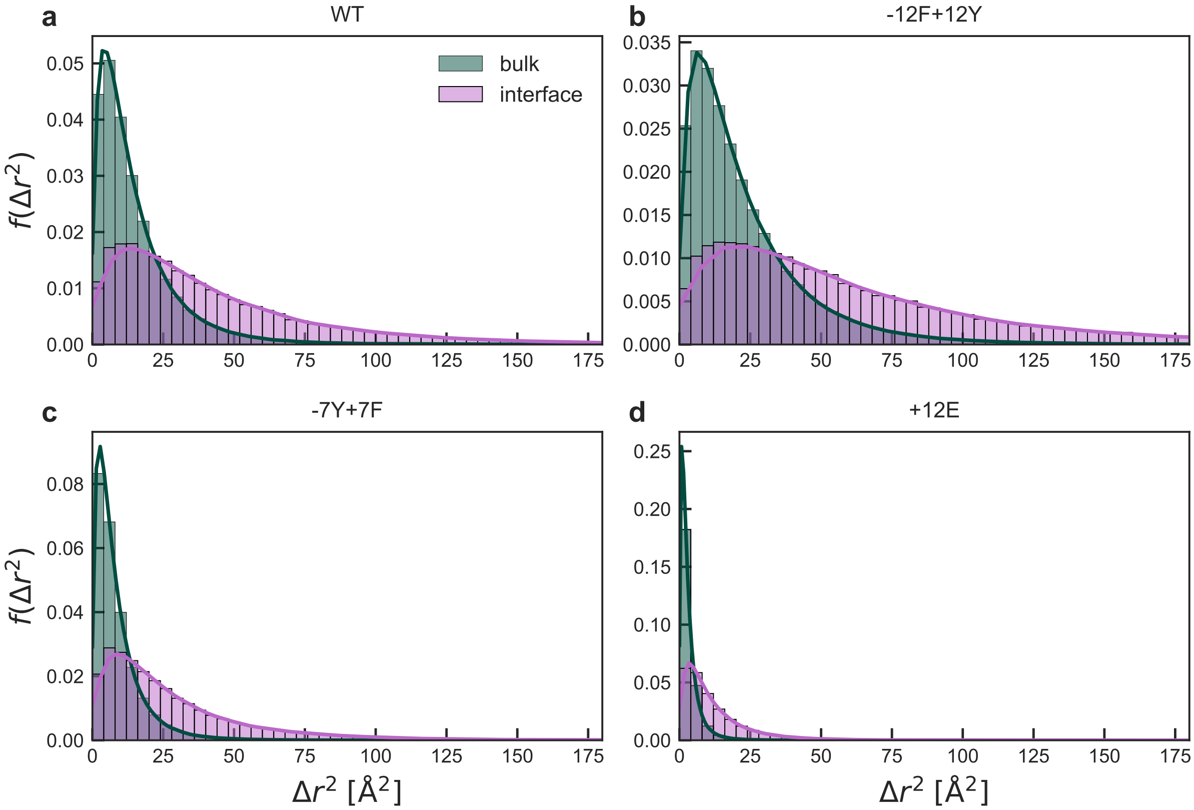


Figure 15: **a**-**d**, PDF of the squared displacement, $\Delta r^2$, computed for $\delta t$ =20 ns at the interface and inside the condensate for WT, -12F+12Y, -7Y+7F, and +12E.

#### Comparison of simulated and Stokes–Einstein diffusion coefficients for single-chain simulations of A1-LCD proteins

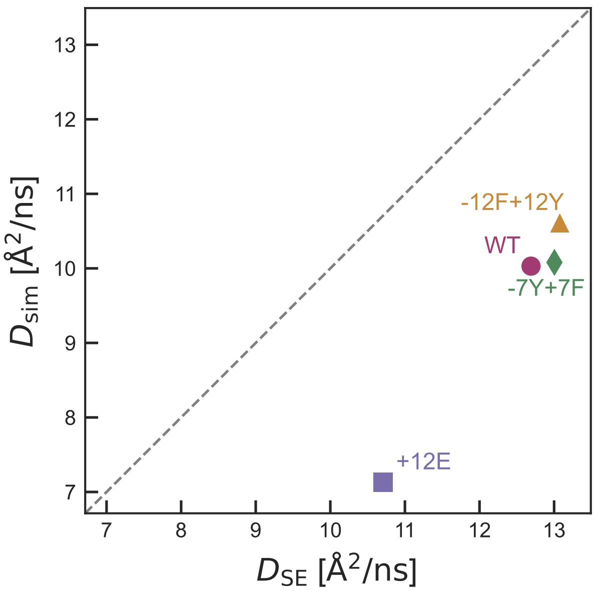


Figure 16: Diffusion coefficients, $D_{\mathrm{sim}}$, obtained from the mean-squared displacement of the protein centre of mass in simulations, and diffusion coefficients estimated using the Stokes–Einstein relation, $D_{\mathrm{SE}}$, for the WT and its variants. The dashed line indicates $D_{\mathrm{sim}} = D_{\mathrm{SE}}$.

To estimate the transnational diffusion for each protein, $D_{\mathrm{sim}}$, we first compute the time-origin averaged mean squared displacement (MSD), $\langle \Delta r^2 \rangle$, of protein's centre of mass within 200 ns trajectory blocks. Then we obtain the diffusion coefficient by a fitting the MSD up to a lag time of 50 ns shown in Fig.6d (main text) with the expression $\langle \Delta r^2 \rangle = 6 D_{\mathrm{sim,PBC}} t$ and apply Yeh–Hummer correction[11,12] accounting for finite-size effects arising from the finite simulation box, resulting in $D_{\mathrm{sim}} = D_{\mathrm{sim,PBC}} + \dfrac{2.837297 k_{\mathrm{B}} T}{6\pi\eta L}$, where $T$ is the absolute temperature, $L$ is the cubic box length and $\eta$ is the shear viscosity of Martini water at the corresponding temperature. The viscosity is determined independently from NVT simulations of Martini water using the Green–Kubo formalism[13]. All single-chain diffusion simulations are performed in the NVT ensemble using the mean box dimensions extracted from previous NPT simulations avoiding possible systematic errors in diffusion coefficients introduced by box-volume fluctuations in NPT simulations[14]. In Fig 16, we compare the estimate diffusion coefficient with diffusion coefficients predicted using the Stokes–Einstein equation[15] $D_{\mathrm{SE}} = \dfrac{k_{\mathrm{B}}T}{6\pi\eta R_{\mathrm{h}}}$, where $k_{\mathrm{B}}$ is the Boltzmann constant, and $R_{\mathrm{h}}$ is protein's hydrodynamic radius estimated through the KirkwoodRiseman equation[16] using internal distances $r_{ij}$ between all pairs of backbone bead $i$ and $j$ as $R_{\mathrm{h}} = \dfrac{1}{\langle r_{ij}^{-1} \rangle_{i \neq j}}$. The theoretical values are of the same order as the ones extracted from simulations, however are larger by 20 % - 30 %. Such deviations are not unexpected, given the approximation of an IDP as a spherical particle in the SE description.
